# Inflammatory markers in pregnancy – surprisingly stable. Mapping trajectories and drivers in four large cohorts

**DOI:** 10.1101/2024.06.19.599718

**Authors:** Frederieke A.J. Gigase, Anna Suleri, Elena Isaevska, Anna-Sophie Rommel, Myrthe G.B.M. Boekhorst, Olga Dmitrichenko, Hanan El Marroun, Eric A.P. Steegers, Manon H.J. Hillegers, Ryan L. Muetzel, Whitney Lieb, Charlotte A.M. Cecil, Victor Pop, Michael Breen, Veerle Bergink, Lot D. de Witte

**Affiliations:** Department of Child and Adolescent Psychiatry, Erasmus MC University Medical Center Rotterdam, Rotterdam, The Netherlands; The Generation R Study Group, Erasmus MC University Medical Center, Rotterdam, The Netherlands; Department of Psychiatry, Icahn School of Medicine at Mount Sinai, New York City, NY, USA; Department of Medical and Clinical Psychology, Tilburg University, Tilburg, The Netherlands; Department of Psychology, Education and Child Studies, Erasmus School of Social and Behavioral Sciences, Erasmus University, Rotterdam, The Netherlands; Department of Obstetrics and Gynecology, Erasmus University Medical Center, Rotterdam, the Netherlands; Department of Radiology and Nuclear Medicine, Erasmus MC University Medical Center, Rotterdam, The Netherlands; Department of Obstetrics, Gynecology and Reproductive Science, Icahn School of Medicine at Mount Sinai, New York City, NY, USA; Department of Epidemiology, Erasmus MC University Medical Center, Rotterdam, The Netherlands; Department of Biomedical Data Sciences, Molecular Epidemiology, Leiden University Medical Center, Leiden, The Netherlands; Department of Psychiatry, Erasmus MC University Medical Centre Rotterdam, Rotterdam, The Netherlands; Department of Human Genetics, Radboud UMC, Nijmegen, The Netherlands; Department of Psychiatry, Radboud UMC, Nijmegen, The Netherlands

**Keywords:** pregnancy, immunology, maternal immune activation, inflammatory marker dynamics, cohort study, intra-individual correlation

## Abstract

Adaptations of the immune system throughout gestation have been proposed as important mechanisms regulating successful pregnancy. Dysregulation of the maternal immune system has been associated with adverse maternal and fetal outcomes. To translate findings from mechanistic preclinical studies to human pregnancies, studies of serum immune markers are the mainstay. The design and interpretation of human biomarker studies require additional insights in the trajectories and drivers of peripheral immune markers. The current study mapped maternal inflammatory markers (C-reactive protein (CRP), interleukin (IL)-1β, IL-6, IL-17A, IL-23, interferon-*γ*) during pregnancy and investigated the impact of demographic, environmental and genetic drivers on maternal inflammatory marker levels in four multi-ethnic and socio-economically diverse population-based cohorts with more than 12,000 pregnant participants. Additionally, pregnancy inflammatory markers were compared to pre-pregnancy levels. Cytokines showed a high correlation with each other, but not with CRP. Inflammatory marker levels showed high variability between individuals, yet high concordance within an individual over time during and pre-pregnancy. Pre-pregnancy body mass index (BMI) explained more than 9.6% of the variance in CRP, but less than 1% of the variance in cytokines. The polygenic score of CRP was the best predictor of variance in CRP (>14.1%). Gestational age and previously identified inflammation drivers, including tobacco use and parity, explained less than 1% of variance in both cytokines and CRP. Our findings corroborate differential underlying regulatory mechanisms of CRP and cytokines and are suggestive of an individual inflammatory marker baseline which is, in part, genetically driven. While prior research has mainly focused on immune marker changes throughout pregnancy, our study suggests that this field could benefit from a focus on intra-individual factors, including metabolic and genetic components.

## Introduction

Adaptations of the immune system during pregnancy have fascinated the immunological field for decades. Immunological changes have likely evolved as a way for placental mammals to tolerate the developing fetus, which expresses both paternal and maternal allo-antigens (1). Accordingly, aberrant adaptations of the maternal immune system during the prenatal period have been suggested to contribute to increased chance of adverse pregnancy and offspring outcomes (2–8). Identifying which factors contribute to atypical immune adaptation during pregnancy is therefore of clinical importance.

Complex immunological processes at the maternal-fetal interface have been mainly studied in rodent models. Preclinical studies have shown that the equilibrium of T-, B- and uterine natural killer (NK) cells (9) and the production of immunosuppressive cytokines by regulatory T cells (10) is important in a typical pregnancy. These models have also shown that a shift in T-helper (Th) cell balance towards increased Th1 cells, complement activation and the depletion of regulatory T cells or NK cells may all lead to adverse pregnancy outcomes, including fetal rejection, impaired fetal growth, and abnormal neurodevelopment (11–16). Mechanistic preclinical models allow for the investigation of the maternal immunological system in ways that cannot be done in humans, including knock-out models to assess the role of particular immune cells, and harvesting of immune system tissues at different timepoints during gestation (17). Together, animal studies have been particularly useful in revealing that a well-balanced immune system is crucial for a successful pregnancy.

In humans, assessment of peripheral blood biomarkers, including cytokines, chemokines and cell-type composition has been employed in an effort to translate preclinical findings to human pregnancy. Signaling molecules such as C-reactive protein (CRP) and cytokines tightly regulate immunological processes and are therefore frequently measured in clinical studies. CRP is an acute-phase protein produced by liver cells and its production is stimulated by proinflammatory cytokines including interleukin (IL)-6 and IL-1. Cytokines are produced by a variety of immune and non-immune cells. During pregnancy, the placenta is an important additional source. IL-1β, IL-6, IL-17A, IL-23 and interferon (IFN)-*γ* have been implicated in early pregnancy processes, such as implantation (18,19) and spiral artery remodeling of the uterine vessels (20). Prior studies have suggested a decrease in cytokine levels from early to mid-pregnancy, reflective of a pro-inflammatory environment in the first trimester followed by an anti-inflammatory state during the second trimester (n=21-707) (21–23). However, a consistent trend throughout pregnancy was not observed across the literature for the majority of cytokines (3). Existing studies were limited in terms of generalizability (i.e. no independent replication cohorts; exclusion of susceptible subgroups) and methodological variability between cohorts (21–23). Large studies with repeated measures, identical assays, and replication cohorts are needed to expand our knowledge of the temporal dynamics of immune markers in pregnancy. To understand the role of atypical immune adaptation, a robust characterization of inflammatory markers, including trajectories and driving forces, throughout normal gestation is warranted.

Well-known triggers such as viral or bacterial infection might be causal to immune activation. Yet, other inflammatory factors such as high body mass index (BMI), tobacco use and parity might also play a role, as suggested by studies of pregnant (21,24–26) and non-pregnant (27) populations. Additionally, the genetic susceptibility to elevated inflammatory marker levels may play a role as indicated for example by a large genome-wide study (GWAS) reporting genetic loci associated with CRP levels (28). While there is evidence that immune dysregulation is associated with adverse maternal and fetal outcomes, it is currently unclear to what extent these factors contribute to dysregulation of the maternal immune system in the general pregnant population. Previous studies have typically employed a cross-sectional design (29), or included single or limited number of biomarkers (30). Identifying robust drivers of systemic immune adaptations will enhance our understanding of successful maintenance of pregnancy, as well as of adverse outcomes.

The two-fold aim of the present study is to i) map inflammatory marker (high sensitivity (HS)-CRP, IL-1β, IL-6, IL-17A, IL-23, IFN-*γ*) patterns throughout pregnancy and ii) investigate the impact of demographic, environmental and genetic factors on maternal inflammatory marker levels. Blood samples were collected at two timepoints in the first half of pregnancy. Maternal serum inflammatory marker levels were measured by multiplex analysis. The Generation R Study, a multi-ethnic population-based prospective pregnancy cohort in the Netherlands (NL) (n=8,082) was used as a discovery cohort (31). Importantly, findings were replicated in three cohorts, namely the Generation C cohort (n=2,535, USA), the Brabant Study cohort (n=587, NL) and the Generation R *Next* Study (n=1,270, NL) with biomarkers assessed at multiple timepoints using identical assays.

## Results

### Characteristics of four cohorts

A total of 8,082 pregnant participants were included in the Discovery Cohort (**Table 1**). Of the included participants, 5,478 participants (67.8%) had repeated cytokine measurements and 5,938 participants (73.5%) had genotype data. In total, 13,467 samples were collected with complete cytokine data. Of these, 13,316 samples also had HS-CRP data. Results were replicated in the Generation C cohort (n=2,535) and the Brabant Study cohort (n=587), hereafter referred to as Replication Cohort I and Replication Cohort II, respectively (**Table 1**). In Replication Cohort I, 3,319 samples were collected from 2,535 participants and 541 of these participants (21.3%) had 2 to 6 repeated cytokine measurements. In Replication Cohort II, 1,170 samples were included from 587 participants and 387 of these participants (65.9%) had 2 to 3 repeated cytokine measurements. The Generation R *Next*, hereafter referred to as the pre-pregnancy cohort, collected samples preconception and during pregnancy. In the pre-pregnancy cohort, 1,779 samples were collected from 1,270 participants and 395 of these participants (31.3%) had up to

**Table 1:**
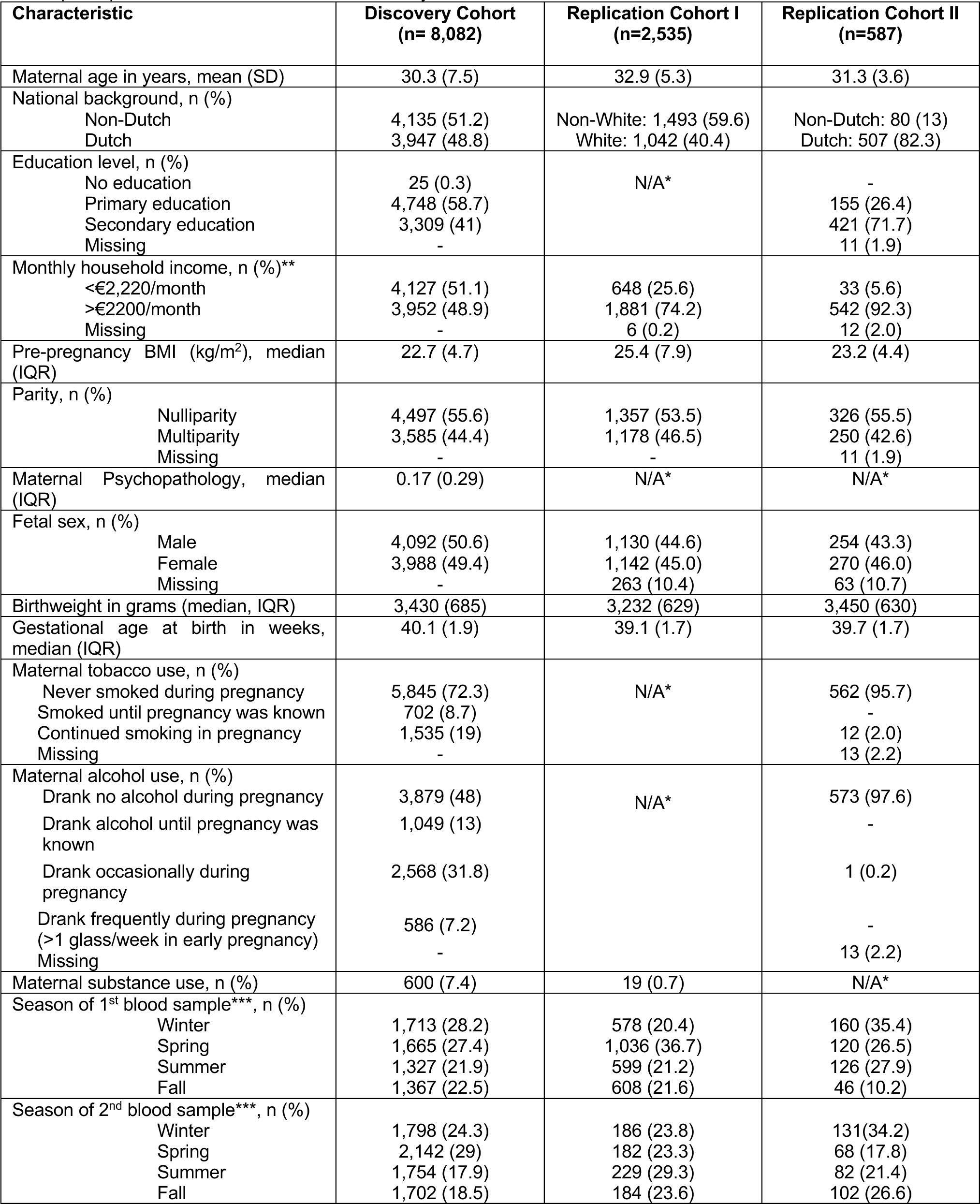
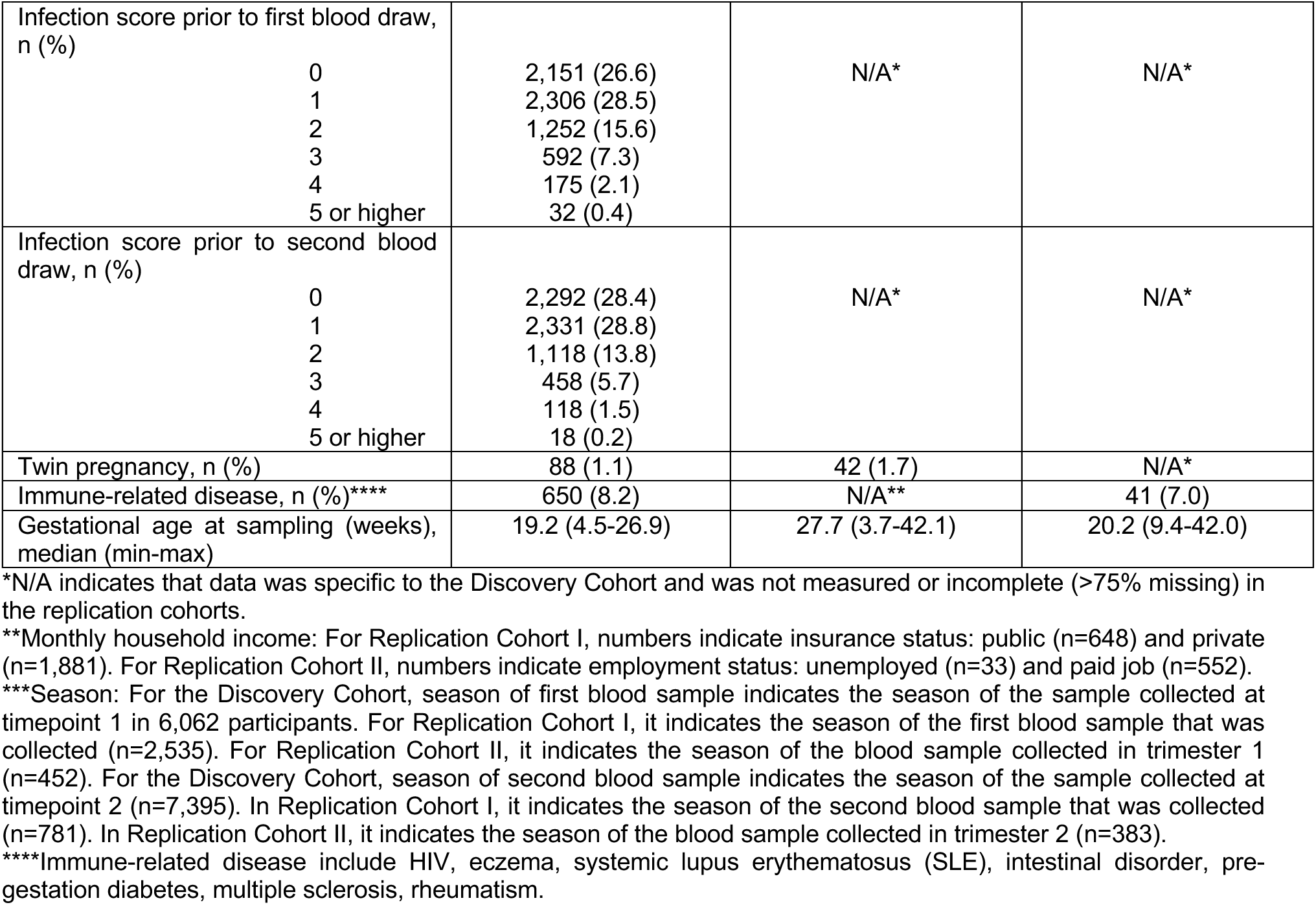
Characteristics of the Discovery Cohort, Replication Cohort I, and Replication Cohort II. A total of 11,204 participants were included in the current study.

5 repeated cytokine measurements preconception and during pregnancy. Study design is visualized in **Figure 1A**. Demographics of the Discovery Cohort are visualized in **Figure 1B-G**.

**Figure 1:**
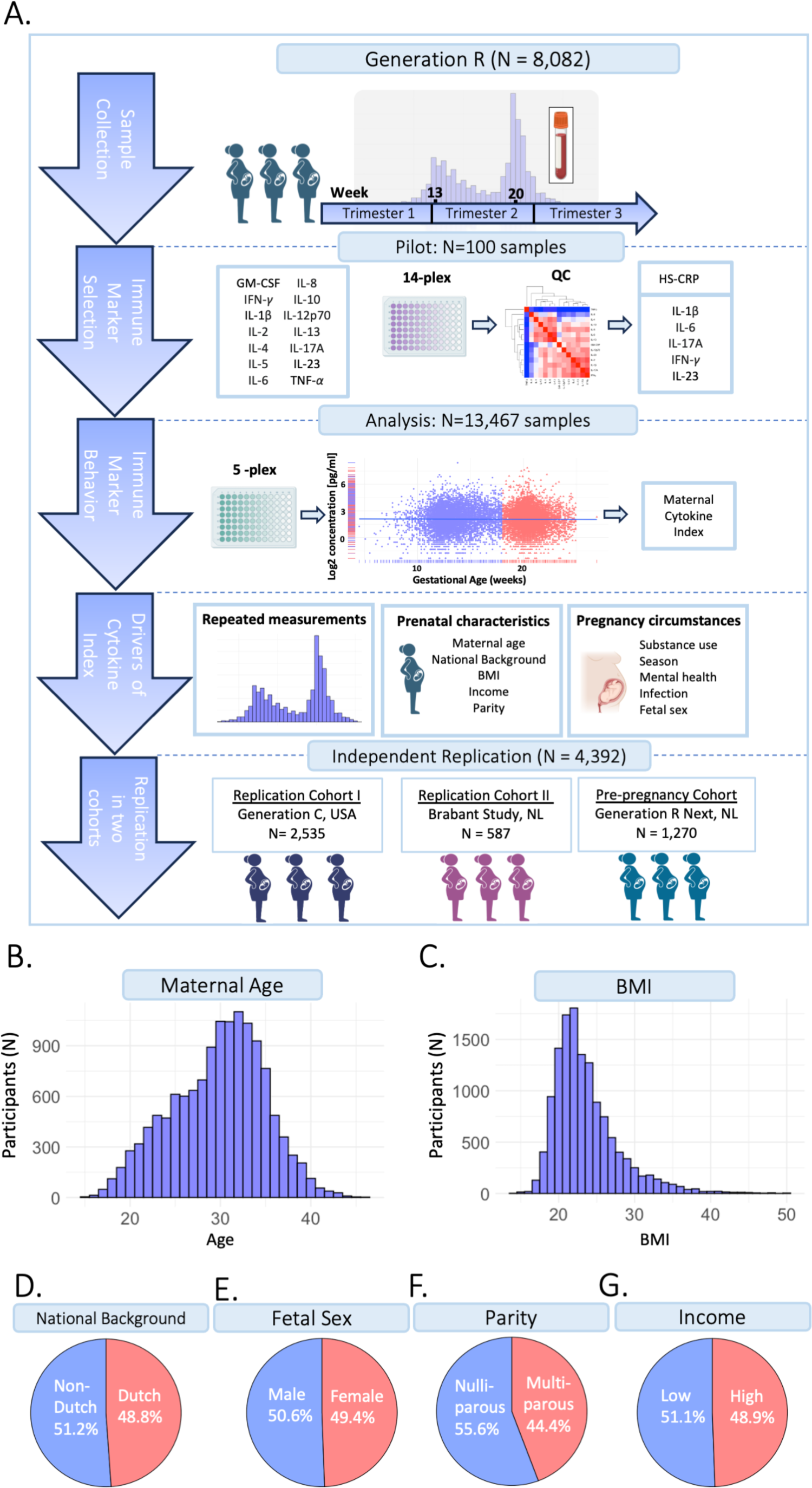
Study design. A) The current study is part of the Generation R Study pregnancy cohort (n=8,082 participants). Sample collection occurred between 2001-2006. Blood samples were collected at two timepoints at median 13.2 weeks (95% range: 9.6-17.6 weeks) and 20.3 weeks (95% range: 18.5-23.3 weeks) gestation, processed and stored for further analyses. A pilot analysis of 14 cytokines was performed in 100 samples. IL-1β, IL-6, IL-17A, IFN-*γ* and IL-23 were selected for analysis in the full cohort, as well as HS-CRP. Participants were excluded based on several exclusion criteria. Data analysis was performed in n=13,467 samples of n=8,082 included participants, of which n=5,478 had a repeated measurement. Inflammatory markers were characterized throughout gestation. A maternal cytokine index was generated for each sample, reflecting inflammatory marker behavior. To identify potential drivers of inflammatory markers, the association between pre-pregnancy characteristics and pregnancy circumstances as predictors of the maternal cytokine index and HS-CRP was assessed. Findings were replicated in two replication cohorts: the Generation C study (n=2,535) and the Brabant Study (n=587) and in a unique pre-pregnancy cohort, the Generation R *Next* study (n=1,270). B) Distribution of maternal age (years). C) Distribution of BMI (kg/m^2^). D) Distribution of national background (Dutch/non-Dutch). E) Distribution of fetal sex (female/male). F) Distribution of parity (nulliparous/multiparous). G) Distribution of household income (low: <€2,220/month/high: >€2200/month).

### Descriptive analysis of maternal inflammatory markers

#### Discovery cohort

In the Discovery Cohort, blood samples were collected at two timepoints during gestation. Median gestational age at sample collection of the first blood sample was 13.2 weeks (95% range: 9.6-17.6 weeks) in early pregnancy, and 20.3 weeks (95% range: 18.5-23.3 weeks) in mid-pregnancy (**Figure 2A**). Of the 13,467 samples, 6,072 (45%) were collected in early pregnancy and 7,395 (55%) in mid pregnancy. Median inflammatory marker levels were 3.95 pg/mL for IL-1β, 1.61 pg/mL for IL-6, 23.98 pg/mL for IL-17A, 1,107.4 pg/mL for IL-23, 14.75 pg/mL for IFN-*γ*, and 4.3 mg/L for HS-CRP (**Table 2**). The univariate analysis showed that group-level inflammatory marker levels were significantly lower at the second timepoint compared to the first (**Figure 2B**; **Table 2**). Repeated measures were median 50 days apart among participants with a repeated measurement (n=5,478 participants, n=10,956 samples). The intra-individual correlation of inflammatory markers between timepoints in these participants was strong (*r*=0.68-0.93, *p*<0.001) (**Figure 2C&D**).

**Figure 2:**
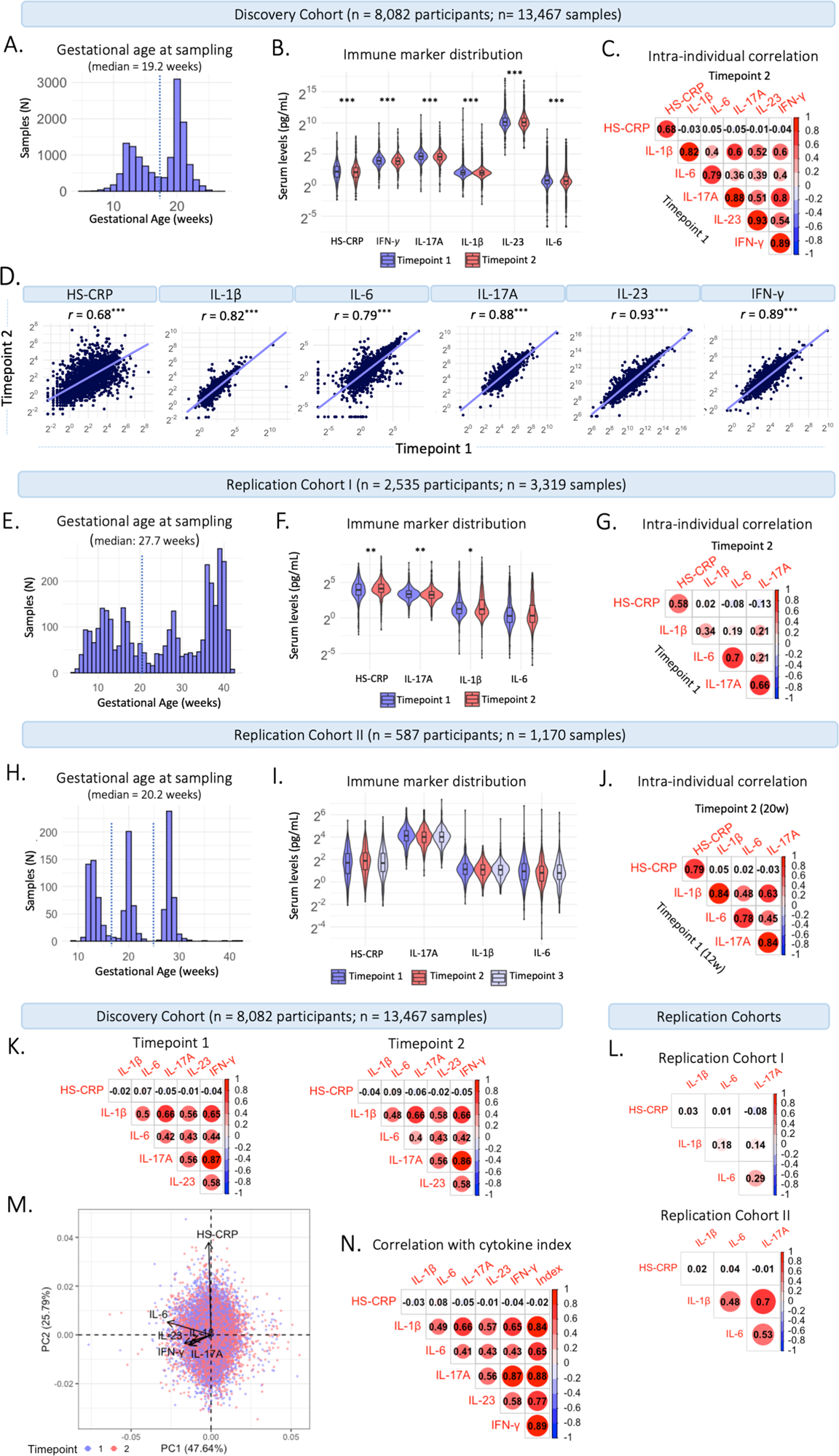
Characteristics of maternal inflammatory markers. A) Distribution of gestational age (weeks) at sample collection in the Discovery Cohort (n=13,467 samples). The dotted line indicates the division between timepoint 1 (trimester 2; 13.2 weeks) and timepoint 2 (trimester 2; 20.3 weeks). B) Violin plots of inflammatory markers measured at timepoint 1 at median 13.2 weeks gestation (95% range: 9.6-17.6 weeks) and timepoint 2 at median 20.3 weeks gestation (95% range: 18.5-23.3 weeks). Cytokines (pg/mL) and HS-CRP (mg/L) are shown on a log2 axis. Univariate analysis at group-level show that inflammatory marker levels are significantly lower at the second timepoint compared to the first (*p*<0.010). C) Intra-individual correlation between samples collected at timepoint 1 and at timepoint 2 among participants with repeated measurements (n=5,478 participants; 10,956 samples). Inflammatory markers were log2 transformed. D) Intra-individual correlation between timepoint 1 (median 13.2 weeks) and timepoint 2 (median 20.3 weeks) in participants with repeated measures in the Discovery Cohort (n=5,478). Correlation coefficients represent Pearson’s r and range from 0.69 to 0.93. Axes are on a log2 scale. E) Distribution of gestational age (weeks) at sample collection in Replication Cohort I (n=3,319 samples). The dotted line indicates the division between timepoint 1 (early gestation; ≤20 weeks) and timepoint 2 (late gestation; >20 weeks). F) Violin plots of inflammatory markers measured in early gestation (≤20 weeks) and late gestation (>20 weeks). Cytokines (pg/mL) and HS-CRP (mg/L) are shown on a log2 axis. Univariate analysis at group-level show that HS-CRP, IL-17A and IL-1β were significantly different between early and late pregnancy (*p*<0.05). G) Intra-individual correlation between the first and second sample collected among participants with repeated measurements (n=541). H) Distribution of gestational age (weeks) at sample collection in Replication Cohort II (n=1,170 samples). The dotted line indicates the division between samples collected at timepoint 1 (trimester 1; 12 weeks), timepoint 2 (trimester 2; 20 weeks), and timepoint 3 (trimester 3; 28 weeks). I) Violin plots of inflammatory markers measured at timepoint 1-3. Cytokines (pg/mL) and HS-CRP (mg/L) are shown on a log2 axis. Univariate analysis at group-level show that inflammatory markers are not different between timepoints. J) Intra-individual correlation between timepoint 1 and 2 among participants with repeated measurements (n=387). K) Correlation between cytokines and HS-CRP at timepoint 1 (left) and timepoint 2 (right) in the Discovery Cohort. L) Correlation between inflammatory markers in Replication Cohort I (top) and Replication Cohort II (bottom). M) Principal component analysis of cytokines and HS-CRP in the Discovery Cohort, indicating high loadings of IL-1β, IL-6, IL-17A, IFN-*γ* and IL-23 on Principal Component (PC) 1 and of HS-CRP on PC 2. Samples are color coded by timepoint. N) Correlation of cytokines and HS-CRP with the maternal cytokine index across all samples in the Discovery Cohort (n=13,467). All correlation coefficients represent Pearson’s r.

**Table 2:** Characteristics of inflammatory markers in the Discovery Cohort. Inflammatory marker levels overall, as well as per timepoint in maternal serum samples obtained from n=8,082 participants in the Discovery Cohort.

| Inflammatory marker | Overall (n=13,467) |  | Timepoint 1* (n=6,062) |  | Timepoint 2* (n=7,395) |  | p-value** |
| --- | --- | --- | --- | --- | --- | --- | --- |
|  | Median (IQR) | Range (min-max) | Median (IQR) | Range (min-max) | Median (IQR) | Range (min-max) |  |
| IL-1β (pg/ml) | 3.95 (2.0) | 0.12 – 5,195.84 | 4.01 (2.03) | 0.28-5,195.84 | 3.91 (1.91) | 0.12-876.15 | <0.001 |
| IL-6 (pg/ml) | 1.61 (1.57) | 0.01 – 507.61 | 1.68 (1.65) | 0.01-507.61 | 1.58 (1.48) | 0.01-168.64 | <0.001 |
| IL-17A (pg/ml) | 23.98 (16.46) | 0.50 – 2,493.46 | 24.64 (16.87) | 0.5-2,493.46 | 23.47 (15.91) | 0.5-1,117.95 | <0.001 |
| IL-23 (pg/ml) | 1107.24 (1019.25) | 29.22 – 143,087.31 | 1,128.63 (1019.76) | 29.22-143,087.31 | 1,088.31 (1017.69) | 63.48-114,077.34 | 0.008 |
| IFN-γ (pg/ml) | 14.75 (10.71) | 0.71 – 999.42 | 15.07 (10.99) | 0.71-999.42 | 14.42 (10.42) | 0.71-512.99 | <0.001 |
|  | Overall (n=13,316) |  | Timepoint 1* (n=5,924) |  | Timepoint 2* (n=7,392) |  |  |
| HS-CRP*** (mg/L) | 4.3 (5.0) | 0.2 – 343.0 | 4.5 (5.7) | 0.2-343.0 | 4.2 (4.6) | 0.2-231.00 | <0.001 |

\*Timepoints according to study design: Timepoint 1 = median 13.2 weeks gestation. Timepoint 2 = median 20.3 weeks gestation.
\*\*Independent samples t-test of normalized cytokines and HS-CRP in early and mid-pregnancy.
\*\*\*HS-CRP was measured in 13,316 samples (5,924 early and 7,392 mid pregnancy samples) from 8,062 participants.

#### Replication Cohort I

In Replication Cohort I, blood samples were collected throughout gestation, at a median gestational age of 27.7 weeks (95% range: 6.9-40.6 weeks) (**Figure 2E**). Median inflammatory marker levels were 2.38 pg/mL for IL-1β, 1.24 pg/mL for IL-6, 9.85 pg/mL for IL-17A, and 16.9 mg/L for HS-CRP (**Supplementary Table 1**). The univariate analysis showed that group-level IL-1β (*p*=0.025) and IL-17A (*p*=<0.001) levels were lower in late pregnancy (>20 weeks), HS-CRP (*p*=<0.001) levels were higher in late pregnancy (>20 weeks) and IL-6 (*p*=0.066) levels were not different between early (<20 weeks) and late (>20 weeks) pregnancy (**Figure 2F; Supplementary Table 1**). Repeated measures were median 48 days apart among participants with a repeated measurement (n=541 participants, n=1,292 samples). The intra-individual correlation of inflammatory markers between early (<20 weeks) and late (>20 weeks) pregnancy in these participants was moderate to strong (*r* =0.34-0.70, *p*<0.001) (**Figure 2G**).

#### Replication Cohort II

In Replication Cohort II, blood samples were collected at 12, 20, and 28 weeks gestation, at a median gestational age of 20.2 weeks (95% range: 12.0-29.5 weeks) (**Figure 2H**). Median inflammatory marker levels were 2.14 pg/mL for IL-1β, 1.79 pg/mL for IL-6, 16.55 pg/mL for IL-17A, and 4.78 mg/L for HS-CRP (**Supplementary Table 2**). The univariate analysis showed that group-level inflammatory marker levels were not significantly different between three timepoints (**Figure 2I, Supplementary Table 2**). Repeated measures were median 50 days apart among participants with a repeated measurement (n=387 participants, n=970 samples). The intra-individual correlation of inflammatory markers between 12 and 20 weeks of pregnancy in these participants was strong (*r* =0.78-0.84, *p*<0.001) (**Figure 2J**) and similar between 12 and 28 weeks (*r* =0.71-0.84, *p*<0.001) and 20 and 28 weeks (*r* =0.76-0.86, *p*<0.001).

#### Construction of the maternal cytokine index

Given that inflammatory markers interact, and to capture systemic inflammatory marker changes, inflammatory markers were summarized in a maternal cytokine index in the Discovery Cohort. Cytokines IL-1β, IL-6, IL-17A, IL-23, and IFN-*γ* correlated moderately to strongly with each other at timepoint 1 and timepoint 2 (*r*=0.40 – 0.86; **Figure 2K**). HS-CRP showed a low correlation with the cytokines at both timepoints (Pearson’s *r*=-0.06 – 0.09). The low correlation of HS-CRP with cytokines was also seen in the two replication cohorts (**Figure 2L**). In a principal component analysis (PCA), cytokines IL-1β, IL-6, IL-17A, IL-23, and IFN-*γ* loaded highly on the first principal component (PC) (84%, 66%, 89%, 77%, 89%, respectively), accounting for 55% of the variance (**Figure 2M**). Given the correlation structure amongst the inflammatory markers and the high loading of HS-CRP on the second PC, HS-CRP was excluded from the maternal cytokine index. The maternal cytokine index correlated strongly with individual cytokines at timepoint 1 (*r*=0.66 – 0.89) and timepoint 2 (*r*=0.64 – 0.89) and correlation between cytokines and the maternal cytokine index was higher compared to correlations among cytokines (**Figure 2N**).

### Aim 1: mapping inflammatory marker dynamics throughout gestation

In the Discovery Cohort, gestational age (range 4.5-26.9 weeks) was significantly associated with HS-CRP (*p*<0.001) and the maternal cytokine index (*p*<0.001) in univariate analyses (**Figure 3A**). HS-CRP showed an increase in early pregnancy, followed by a gradual decrease (**Figure 3A**). The maternal cytokine index showed a gradual decrease throughout early and mid-pregnancy (**Figure 3A**). Individual cytokines displayed similar trajectories across gestational age (**Figure 3A**). In Replication Cohort I, gestational age at sample collection (range 3.7-42.1 weeks) was significantly associated with HS-CRP, IL-1β, IL-6, and IL-17A (*p*<0.001) in univariate analyses (**Figure 3B**). Similar to the Discovery Cohort, HS-CRP showed an increase in early pregnancy, followed by a gradual decrease. Cytokines showed a gradual decrease throughout early pregnancy, followed by an increase of IL-1β and IL-6, but not IL-17A, in the third trimester. In Replication Cohort II, gestational age at sample collection (range 9.4-42.0 weeks) was not significantly associated with inflammatory markers (HS-CRP (*p*=0.192), IL-1β (*p=*0.372), IL-6 (*p*=0.111), and IL-17A (*p*=0.102)) in univariate analyses (**Figure 3C**).

**Figure 3.**
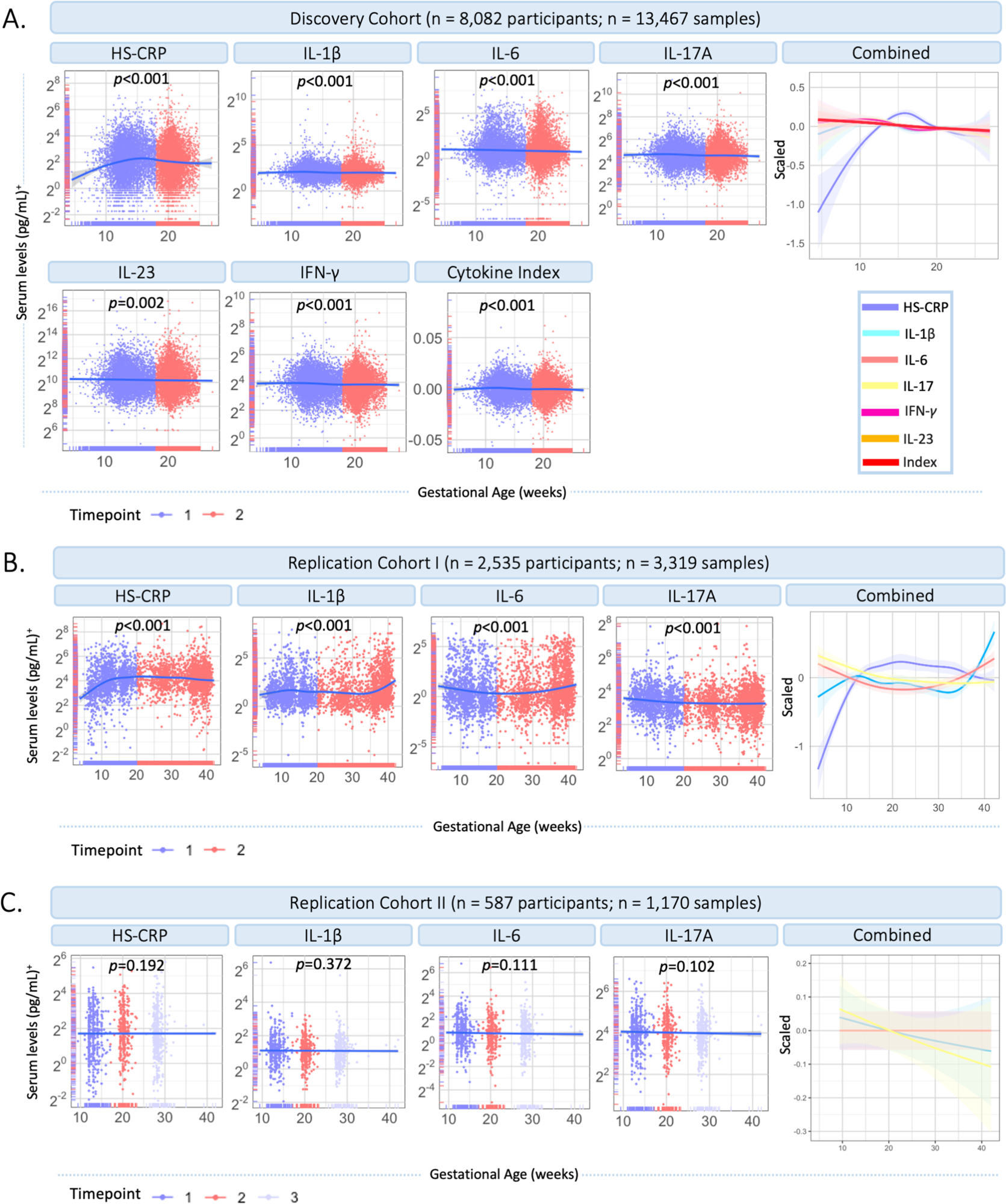
Inflammatory marker dynamics throughout gestation. A) Inflammatory markers across gestational age in the Discovery Cohort (n=13,467 samples). Measurements are color coded by timepoint according to study design (timepoint 1 = median 13.2 weeks; timepoint 2 = 20.3 weeks). Inflammatory markers and the maternal cytokine index are scaled and combined in one plot (right). B) Inflammatory markers across gestational age in Replication Cohort I (n=3,319 samples). Measurements are color coded by timepoint according to study design (timepoint 1 = ≤20 weeks gestation; timepoint 2 = >20 weeks gestation). Inflammatory markers are scaled and combined in one plot (right). C) Inflammatory markers across gestational age in Replication Cohort II (n=1,170 samples). Measurements are color coded by timepoint according to study design (timepoint 1 = median 12 weeks; timepoint 2 = median 20 weeks; timepoint 3 = median 28 weeks). Inflammatory markers are scaled and combined in one plot (right). <u>Legend:</u> A general additive modeling (GAM) line was fitted for each marker indicated with the blue line and a 95% confidence interval on a log2 y-axis. P-values indicate significance of the non-linear association with gestational age. Color coding of inflammatory markers is identical in figure A-C. ^+^HS-CRP was measured in mg/L, cytokines were measured in pg/ml.

### Aim 2: Drivers of maternal inflammatory markers

#### Environmental and genetic drivers of HS-CRP and the maternal cytokine index (univariate analyses)

To characterize drivers of the maternal inflammatory marker landscape, we assessed the association of multiple predictors with HS-CRP and the maternal cytokine index. In the Discovery Cohort (**Figure 4A-C; Supplementary Table 3**), most of the variance in HS-CRP (55.6%) and the maternal cytokine index (87.4%) is explained by within-individual effects. In addition, the variance in HS-CRP levels was partly driven by pre-pregnancy BMI (9.6%) (**Figure 4A-C; Supplementary Table 3**). Less than 1% of the variance in inflammatory markers was explained by pre-pregnancy characteristics and pregnancy circumstances (**Figure 4A-C**). These findings were replicated in both replication cohorts as the majority of the variance in HS-CRP (Replication Cohort I = 33.4%; Replication Cohort II = 58.1%) and cytokines (Replication Cohort I = 26.3-71.5%; Replication Cohort II = 74.7-84.2) was explained by within-individual effects (**Figure 4D; Supplementary Table 3**). Pre-pregnancy BMI contributed to the variance in HS-CRP in both Replication Cohort I (14.9%) and Replication Cohort II (16.1%) (**Figure 4D; Supplementary Table 3**). Less than 1% of the variance in inflammatory markers was explained by pre-pregnancy characteristics and pregnancy circumstances (**Figure 4D**). Across all cohorts, the remaining variance was attributed to residual factors (Discovery Cohort = 6.6%-32.1%; Replication Cohort I = 28%-72.9%; Replication Cohort II = 14.7%-25.1%) (**Figure 4A-D; Supplementary Table 3)**. Across all cohorts, HS-CRP and pre-pregnancy BMI correlated moderately (*r*= 0.32 – 0.41) (**Figure 4E**). Based on the high intra-individual correlation of inflammatory markers and the lack of variance explained by pre-pregnancy characteristics and pregnancy circumstances, we further investigated whether HS-CRP levels were genetically determined. We constructed a PGS of CRP in all ancestries and European ancestries in the discovery cohort, both of which showed a moderate correlation (*r* =0.4) with serum HS-CRP levels (**Figure 5A**). Of the variance in HS-CRP, 14.1% was explained by the CRP PGS in all ancestries and 15.7% was explained by the CRP PGS in European ancestries (**Figure 5B**).

**Figure 4.**
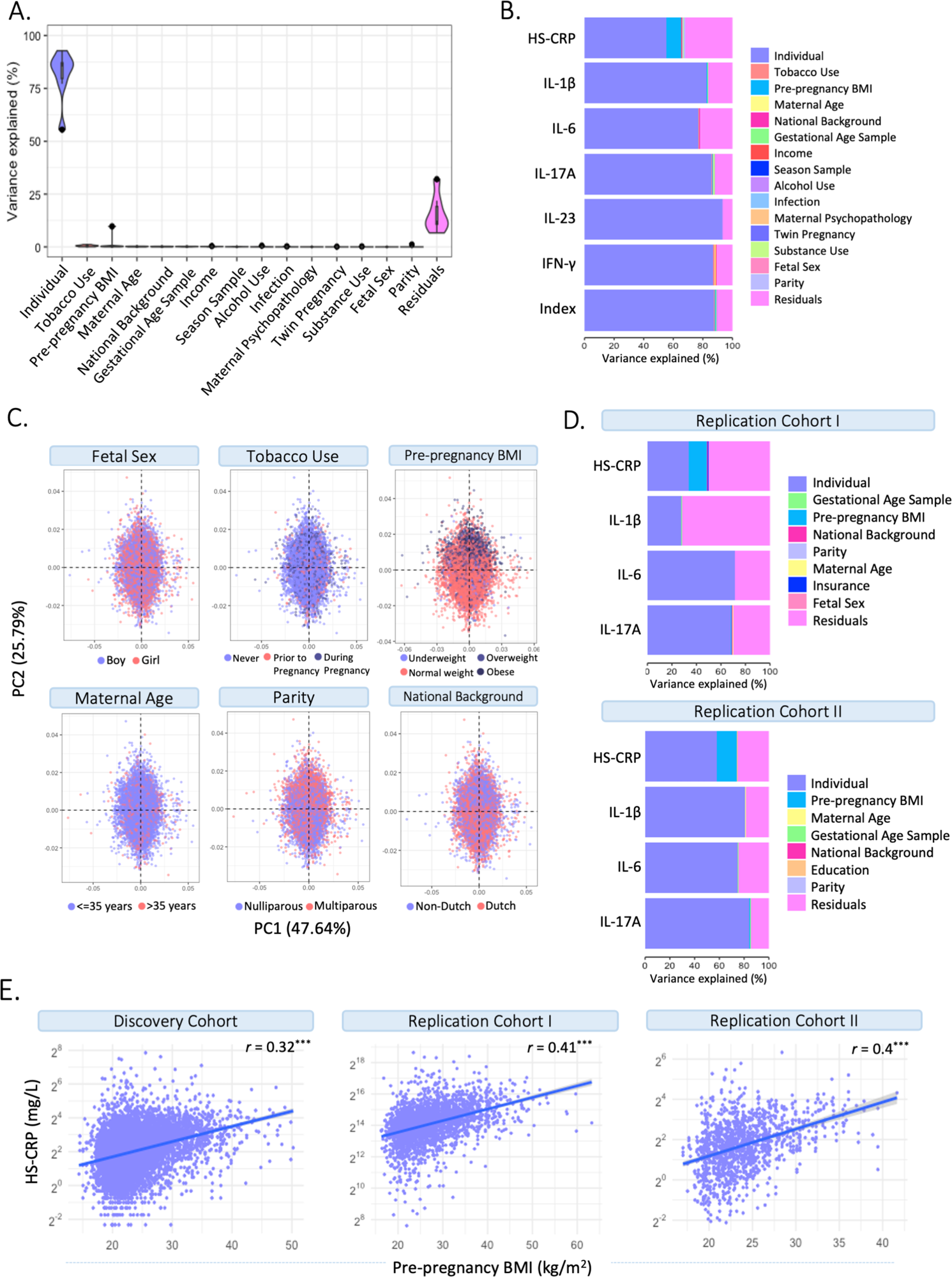
Identifying drivers of maternal inflammatory markers. A) Variance partitioning analysis revealed the variance explained by potential drivers in HS-CRP, cytokines, and the maternal cytokine index. B) The majority of the variance in HS-CRP (55.6%) and the maternal cytokine index (87.4%) is explained by factors captured within the individual. The variance in HS-CRP levels was partly driven by pre-pregnancy BMI (9.6%). The remaining variance was attributed to residual factors (6.6-32.1%). Other factors explained <1% variance in inflammatory markers. C) Distribution of potential drivers of inflammatory markers across Principal Component Analysis (PCA). D) Variance partitioning analysis revealed the variance explained by potential drivers in inflammatory markers in Replication Cohort I (left) and Replication Cohort II (right). The majority of variance in HS-CRP (Replication Cohort I = 33.4%; Replication Cohort II = 58.1%) and cytokines (Replication Cohort I = 26.3-71.5%; Replication Cohort II = 74.7-84.2) was explained by factors captured within the individual. Pre-pregnancy BMI contributed to the variance in HS-CRP (Replication Cohort I=14.9%; Replication Cohort II=16.1%). The remaining variance was attributed to residual factors (Replication Cohort I = 28%-72.9%; Replication Cohort II = 14.7%-25.1%). E) HS-CRP and pre-pregnancy BMI show a moderate correlation (*r* = 0.32) in the Discovery Cohort and both Replication Cohort I (*r* = 0.41) and Replication Cohort II (*r* = 0.4).

**Figure 5.**
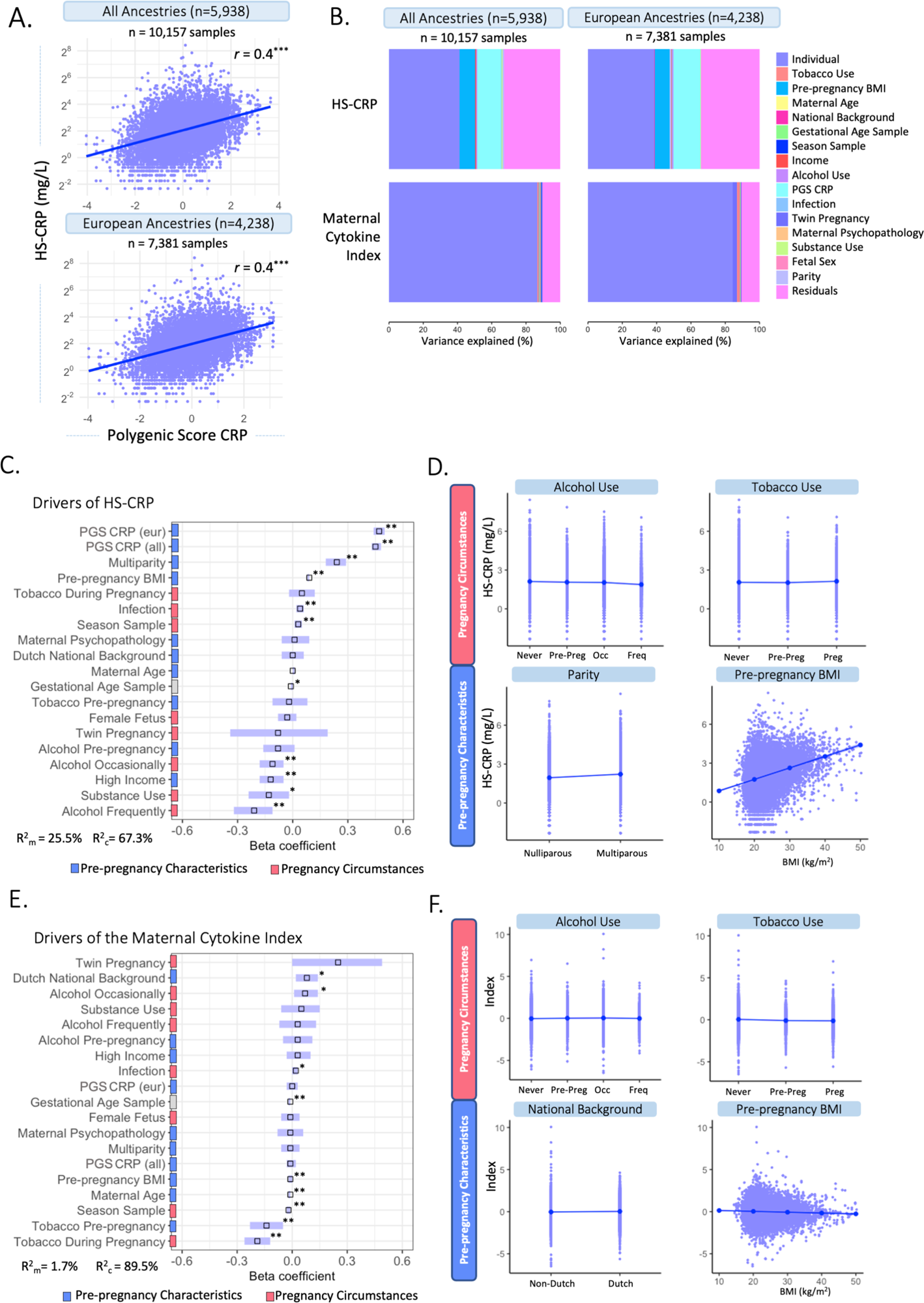
Multivariable associations between potential drivers and HS-CRP and the maternal cytokine index. A) A polygenic Score (PGS) was computed for CRP among participants of all ancestries (n=5,938) and participants of European ancestry (n=4,238). CRP PGS were correlated with HS-CRP levels. Y-axis is on a log2 scale. B) Variance explained by potential drivers in HS-CRP and the maternal cytokine index, including the PGS of all ancestries (left) and European ancestries (right). C) Mixed effects linear regression model of potential drivers and their association with HS-CRP was performed among participants with a CRP PGS (n=5,938 participants, n=10,157 samples). Forest plot shows beta coefficients and 95% confidence interval of selected predictors with HS-CRP. D) Visualization of the association of various pre-pregnancy characteristics and pregnancy circumstances with HS-CRP. E) Mixed effects linear regression model of potential drivers and their association with the maternal cytokine index was performed among participants with a CRP PGS(n=5,938 participants, n=10,157 samples). Forest plot shows beta coefficients and 95% confidence interval of selected predictor. F) The association between various pre-pregnancy characteristics and pregnancy circumstances with the maternal cytokine index. <u>Legend:</u> C-F: Pre-pregnancy characteristics are indicated by blue; pregnancy circumstances are indicated by red. C&E: R^2^_m_ indicates the marginal R squared of the mixed effects regression model, R^2^_c_ indicates the conditional R-squared of the mixed effects regression model after adding individual as a random intercept. Asterisks indicate significance level after multiple testing correction (*p<0.05, **p<0.01, ***p<0.001). D&F: Alcohol categories are ‘never drank’, ‘continued drinking until pregnancy was known (pre-preg)’, ‘drank occasionally (occ)’ and ‘drank frequently (freq)’. Tobacco categories are ‘never smoked’, ‘continued tobacco use until pregnancy was known (pre-preg)’ and ‘continued tobacco use during pregnancy (preg)’.

#### Environmental and genetic drivers of HS-CRP (multivariable analyses)

Mixed effects logistic regression models were performed to identify potential drivers of HS-CRP among participants with a CRP PGS (n=5,938 participants, n=10,157 samples). The CRP PGS was the strongest driver of HS-CRP among participants of all ancestry (β=0.45, 95% CI = 0.43; 0.48, p=0.003) and among participants of European ancestry (β=0.47, 95% CI = 0.44; 0.50, p=0.003) (**Figure 5C**). In addition, we found significant associations with pre-pregnancy characteristics BMI (β=0.09, 95% CI = 0.08; 0.09, p=0.003), high household income (β =-0.12, 95% CI =-0.18; -0.05, p=0.003), and multiparity (β =0.24, 95% CI = 0.18-0.29, p=0.003), as well as pregnancy circumstances, namely occasional and frequent alcohol use during pregnancy (β =-0.11, 95% CI = -0.18; -0.05, p= 0.003 and β =-0.21, 95% CI = -0.32; -0.11, p=0.003, respectively), season at first blood draw (β =0.03, 95% CI = 0.01; 0.05, p= 0.010) and infection score during pregnancy (β =0.04, 95% CI = 0.02; 0.06, p=0.003) in the Discovery Cohort (**Figure 5C-D**, **Table 3**). Maternal age (β =0.00, 95% CI = 0.00; 0.01, p=0.470), national background (β =0.00, 95% CI = -0.06; 0.06, p=0.945), maternal psychopathology (β =0.01, 95% CI = -0.06; 0.09, p=0.815), female fetus (β =-0.03, 95% CI = -0.08; 0.02, p=0.358), and twin pregnancy (β =-0.08, 95% CI = -0.34; 0.19, p=0.678) were not associated with HS-CRP (**Figure 5C-D**, **Table 3**). The marginal variance (i.e., variance explained by fixed effects) was 25.5%. Taking into account the individual, modeled as a random intercept, the conditional variance explained was 67.3%. In replication cohorts, pre-pregnancy BMI was significantly associated with increased HS-CRP levels in Replication Cohort I (β = 0.07, 95% CI = 0.06; 0.07, p=0.004) and Replication Cohort II (β = 0.14, 95% CI = 0.11; 0.16, p=0.009). White ethnicity was associated with decreased HS-CRP levels in Replication Cohort I (β = -0.14, 95% CI = -0.24; -0.05, p=0.005), but this was not the case for national background in Replication Cohort II (β = 0.15, 95% CI = -0.16; 0.46, p=0.623). Other maternal factors showed no significant associations with HS-CRP in Replication Cohort I and II (**Supplementary Figure 4B-C, Supplementary Table 4**).

**Table 3:** Mixed effects models to investigate the association between selected predictors and HS-CRP and the maternal cytokine index. Analyses were performed among participants with an available polygenic score of CRP (n=5,938) and include n=10,157 samples.

|  | HS-CRP (n=10,157) |  |  |  | Maternal cytokine index* (n=10,157) |  |  |  |
| --- | --- | --- | --- | --- | --- | --- | --- | --- |
| | $\beta$ | 95% CI | P-value | Corrected P-value* | $\beta$ | 95% CI | P-value | Corrected P-value* |
| Maternal age | 0.00 | 0.00 – 0.01 | 0.371 | 0.470 | -0.01 | -0.02 – -0.01 | **<0.001** | **0.005** |
| Dutch national background | 0.00 | -0.06 – 0.06 | 0.945 | 0.945 | 0.08 | 0.02 – 0.14 | **0.006** | **0.014** |
| Pre-pregnancy BMI | 0.09 | 0.08 – 0.09 | **<0.001** | **0.003** | -0.01 | -0.02 – 0.00 | **0.003** | **0.010** |
| PGS CRP (all ancestry) | 0.45 | 0.43 – 0.48 | **<0.001** | **0.003** | -0.01 | -0.03 – 0.02 | 0.477 | 0.604 |
| PGS CRP (European ancestry)** | 0.47 | 0.44 – 0.50 | **<0.001** | **0.004** | 0.00 | -0.03 – 0.03 | 0.840 | 0.887 |
| High income | -0.12 | -0.18 – -0.05 | **<0.001** | **0.003** | 0.03 | -0.03 – 0.10 | 0.284 | 0.450 |
| Multiparity | 0.24 | 0.18 – 0.29 | **<0.001** | **0.003** | -0.01 | -0.06 – 0.04 | 0.681 | 0.719 |
| Maternal Psychopathology | 0.01 | -0.06 – 0.09 | 0.772 | 0.815 | -0.01 | -0.08 – 0.06 | 0.835 | 0.835 |
| Maternal tobacco use***<br>Pre-pregnancy | -0.02 | -0.11 – 0.08 | 0.736 | 0.815 | -0.14 | -0.23 – -0.05 | **0.002** | **0.008** |
| Maternal tobacco use***<br>During pregnancy | 0.05 | -0.02 – 0.12 | 0.198 | 0.289 | -0.19 | -0.26 – -0.12 | **<0.001** | **0.005** |
| Maternal alcohol use***<br>Pre-pregnancy | -0.08 | -0.16 – 0.01 | 0.072 | 0.124 | 0.03 | -0.05 – 0.11 | 0.470 | 0.604 |
| Maternal alcohol use***<br>Occasionally during pregnancy | -0.11 | -0.18 – -0.05 | **0.001** | **0.003** | 0.07 | 0.01 – 0.14 | **0.019** | **0.040** |
| Maternal alcohol use***<br>Frequently during pregnancy | -0.21 | -0.32 – -0.11 | **<0.001** | **0.003** | 0.03 | -0.07 – 0.13 | 0.560 | 0.665 |
| Substance use during pregnancy | -0.13 | -0.24 – -0.02 | **0.018** | **0.034** | 0.05 | -0.06 – 0.15 | 0.381 | 0.557 |
| Female fetus pregnancy | -0.03 | -0.08 – 0.02 | 0.264 | 0.358 | -0.01 | -0.06 – 0.04 | 0.611 | 0.683 |
| Twin pregnancy | -0.08 | -0.34 – 0.19 | 0.571 | 0.678 | 0.25 | 0.00 – 0.49 | 0.050 | 0.088 |
| Season Sample | 0.03 | 0.01 – 0.05 | **0.004** | **0.010** | -0.02 | -0.03 – -0.01 | **0.001** | **0.005** |
| Infection score** | 0.04 | 0.02 – 0.06 | **<0.001** | **0.003** | 0.02 | 0.00 – 0.03 | **0.006** | **0.014** |
| Gestational age of the sample | -0.01 | -0.01 – 0.00 | **0.010** | **0.021** | -0.01 | -0.02 – -0.01 | **<0.001** | **0.005** |
| Cytokine batch | 0.00 | 0.00 – 0.00 | 0.196 | 0.289 | 0.00 | 0.00 – 0.00 | 0.051 | 0.088 |
| Marginal R<sup>2</sup> / Conditional R<sup>2</sup> | 0.255 / 0.673 |  |  |  | 0.017/0.895 |  |  |  |

+ Maternal cytokine index was scaled
\*p-value after Benjamini-Hochberg correction.
\*\*Interpreted in separate model.
\*\*\* Reference group: Never smoked during pregnancy/Never drank during pregnancy

#### Environmental and genetic drivers of the maternal cytokine index (multivariable analyses)

Next, we identified several significant associations between potential drivers of inflammation and the maternal cytokine index, including pre-pregnancy characteristics such as maternal age (β =-0.01, 95% CI = -0.02; -0.01, p=0.005), pre-pregnancy BMI (β =-0.01, 95% CI = -0.02; 0.00, p=0.003), and Dutch national background (β=0.08, 95% CI = 0.02; 0.14, p=0.014), as well as pregnancy circumstances including tobacco use pre-pregnancy and during pregnancy (β =-0.14, 95% CI = -0.23; -0.05, p=0.008; β =-0.19, 95% CI = -0.26; -0.12, p=0.005, respectively), occasional alcohol use during pregnancy (β =0.07, 95% CI = 0.01; 0.14, p= 0.040), season at first blood draw (β =-0.02, 95% CI = -0.03; -0.01, p=0.005), and infection score during pregnancy (β =0.02, 95% CI = 0.00 – 0.03, p= 0.014) in the Discovery Cohort (**Figure 5E-F**, **Table 3**). Multiparity (β = -0.01, 95% CI = -0.06; 0.04, p=0.719), maternal psychopathology (β = -0.01, 95% CI = -0.08; 0.06, p=0.835), substance use during pregnancy (β = 0.05, 95% CI = -0.06; 0.15, p=0.557), female fetus (β = -0.01, 95% CI = -0.06; 0.04, p=0.683), twin pregnancy (β = 0.25, 95% CI = 0.00; 0.49, p=0.088), and the CRP PGS for all ancestry (β = -0.01, 95% CI = -0.03; 0.02, p=0.604) and for European ancestry (β = 0.00, 95% CI = -0.03; 0.03, p=0.887) were not associated with the maternal cytokine index (**Figure 5E-F**, **Table 3**). The marginal variance (i.e., variance explained by fixed effects) was 1.7%. Taking into account the individual, modeled as a random intercept, the conditional variance explained was 89.5%.

#### Comparison pre-pregnancy and pregnancy

The contribution of the PRS of CRP to serum levels of HS-CRP, suggested a genetic contribution to inflammatory marker variance during pregnancy. Together with the high intra-individual correlation of cytokines, this led to the hypothesis that immune markers might not show major changes during pregnancy compared to preconception. We assessed HS-CRP and cytokine behavior in a pre-pregnancy cohort, the Generation R *Next* study. Samples were collected preconceptionally at median 11 weeks prior to conception (95% range: 74.5-0 weeks prior to conception, n=676), and during pregnancy at median 8.4 weeks gestation (95% range: 6.4-12.9 weeks, n=1,103) (**Figure 6A**). Median inflammatory marker levels were 4.82 pg/mL for IL-1β, 1.92 pg/mL for IL-6, 59.47 pg/mL for IL-17A, 1,209.5 pg/mL for IL-23, 18.89 pg/mL for IFN-*γ*, and 1.4 mg/L for HS-CRP (**Supplementary Table 5**). The univariate analysis showed that inflammatory marker levels were not significantly different between preconception and first trimester for IL-1β, IL-6, IL-17A, IL-23 and IFN-*γ* (**Supplementary Table 5; Figure 6B**). HS-CRP was significantly higher in the first trimester compared to preconception (**Supplementary Table 5; Figure 6B**). The intra-individual correlation of cytokines preconception and during pregnancy was strong (r=0.79-0.96, *p*<0.001) (**Figure 6C**). HS-CRP showed a moderate correlation between preconception and first trimester samples (r=0.45, <0.001) (**Figure 6C**). In the pre-pregnancy cohort, gestational age at sample collection (range 85 weeks prior to preconception - 17.6 weeks gestation) was significantly associated with HS-CRP (p<0.001), but not with IL-1β (p=0.360), IL-6 (p=0.530), IL-17A (0.313), IL-23 (p=0.376) and IFN-*γ* (p=0.110) in univariate analyses (**Figure 6D&E**). Similar to the Discovery Cohort, HS-CRP showed an increase in early pregnancy.

**Figure 6.**
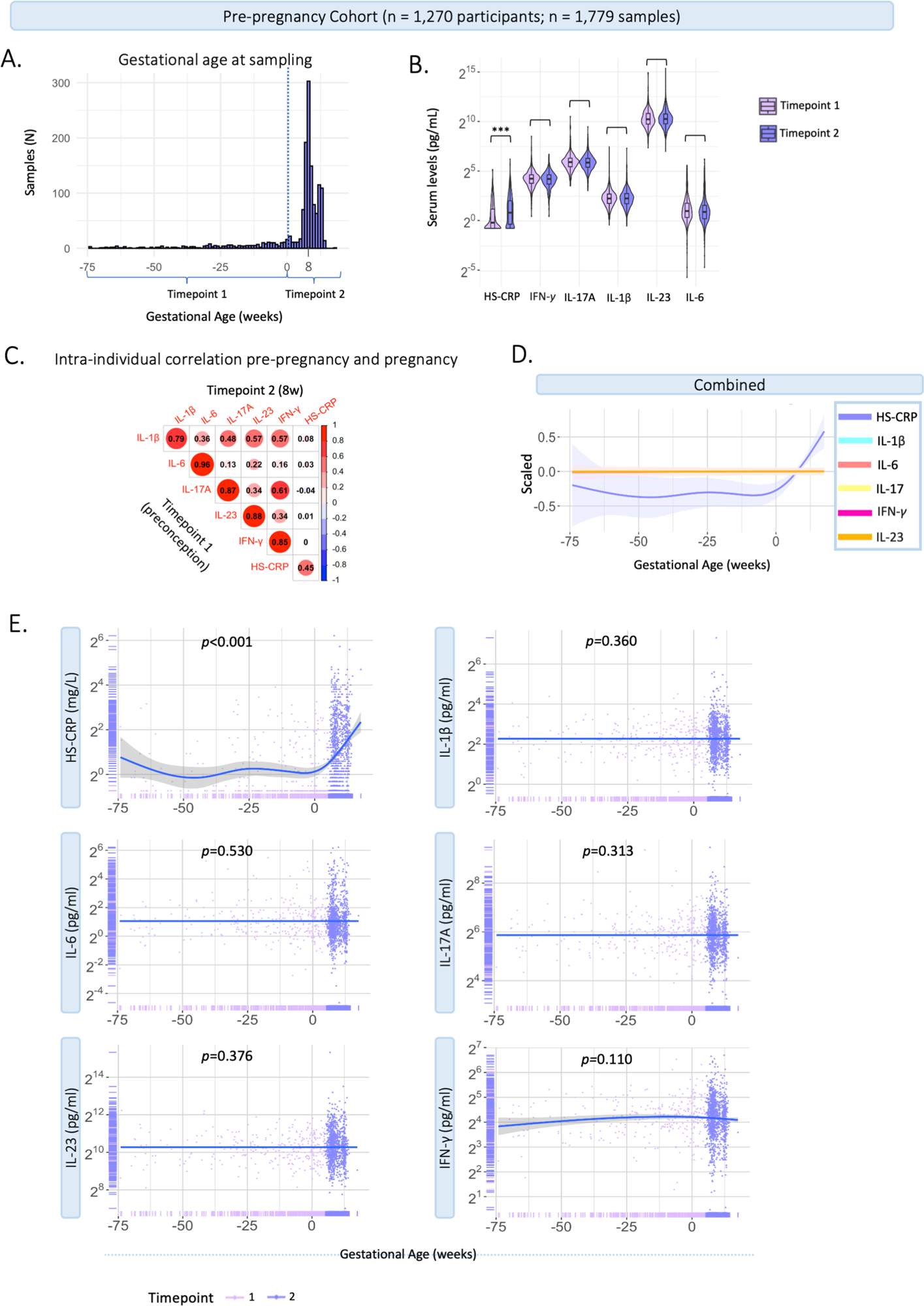
Characteristics of inflammatory markers pre-pregnancy and during pregnancy. Generation R *Next* cohort. A) Distribution of gestational age (weeks) at sample collection (n=1,779 samples). The dotted line indicates the distinction between timepoint 1 (preconception samples), and timepoint 2 (trimester 1; median 8.4 weeks gestation). B) Violin plots of inflammatory markers measured at timepoint 1 (preconceptionally; median 11.4 weeks before conception) and timepoint 2 (median 8.4 weeks gestation (95% range: 6.4-12.9 weeks). Cytokines (pg/mL) and HS-CRP (mg/L) are shown on a log^2^ axis. Univariate analysis at group-level show that cytokine levels were not significantly different between the first trimester and preconception. HS-CRP was significantly higher in the first trimester compared to preconception. C) Intra-individual correlation between samples collected preconception (timepoint 1) and during pregnancy (timepoint 2) among participants with repeated measurements (n=395). Inflammatory markers were log2 transformed. Pearson correlation is shown. D) Inflammatory markers are scaled and combined in one plot. E) Inflammatory markers preconception and during pregnancy in the pre-pregnancy cohort (n=1,779 samples). Measurements are color coded by timepoint according to study design. A general additive modeling (GAM) line was fitted for each marker indicated with the blue line and a 95% confidence interval on a log2 y-axis. P-values indicate significance of the non-linear association with gestational age. HS-CRP was measured in mg/L, cytokines were measured in pg/ml.

#### Sensitivity analyses

The PGS of CRP remained the largest predictor of HS-CRP in several sensitivity analyses in the Discovery Cohort, namely: i) excluding 163 samples (n=123 participants) with potential human anti-animal antibodies (HAAA), which can introduce technical interference in immunological assays (**Supplementary Figure 2A-B, Supplementary Table 6**); ii) excluding 855 samples of 495 participants with immune-related diseases (HIV, eczema, systemic lupus erythematosus (SLE), intestinal disorder, pre-gestation diabetes, multiple sclerosis, rheumatism) (**Supplementary Figure 2C-D, Supplementary Table 7**); and iii) excluding 174 outlier samples (n=147 participants) (**Supplementary Figure 2E-F, Supplementary Table 8**). Additionally, a sensitivity analysis was performed using the delta (change) between the two timepoints for HS-CRP and the maternal cytokine index. Infection score in early pregnancy (*r* =0.08, *p*<0.01) was correlated with the delta HS-CRP, and season (*r* =0.12, *p*=0.04) was correlated with the delta maternal cytokine index. Correlation coefficients between predictors and the HS-CRP delta (*r* =0.00-0.08) and maternal cytokine index delta (*r* =0.00-0.12) were low (**Supplementary Figure 3A)**. The proportion of participants reporting an infection was significantly higher in the high-low HS-CRP group compared to the stable and low-high HS-CRP groups (**Supplementary Figure 3B**). Moreover, parity, season of first blood draw, and tobacco use during pregnancy significantly differed across categories of delta HS-CRP (**Supplementary Figure 3B**). Among maternal cytokine index delta categories, season of first blood draw differed significantly (**Supplementary Figure 3C**). A sensitivity analysis including only participants with a high absolute delta HS-CRP (5^th^ and 95^th^ quantiles) (n=435 participants; n=854 samples) no longer revealed significant associations between important drivers including multiparity, pre-pregnancy BMI, alcohol use, and household income with HS-CRP (**Supplementary Figure 2G-H, Supplementary Table 9)**. The associations between predictors and individual cytokines in the Discovery Cohort are shown in **Supplementary Figure 4A**. White ethnicity, BMI, and maternal age were associated with IL-17A in Replication Cohort I. No significant associations of potential drivers were found with IL-1β and IL-6 in both Replication Cohorts (**Supplementary Figure 4B-C**).

## Discussion

This study is the largest investigation to date of inflammatory markers during pregnancy in four multi-ethnic and socio-economically diverse pregnancy cohorts (>12,000 participants, The Netherlands and USA). Repeated measures enabled the thorough characterization of inflammatory marker trajectories and the assessment of prominent inflammation drivers. Across cohorts, we consistently found strong correlations among cytokines, but no association between cytokines and HS-CRP. Results revealed a high intra-individual correlation of inflammatory markers measured at different time points during gestation as well as preconception using a unique pre-pregnancy cohort. Although we identified gestational age-dependent changes, results showed that gestational age at sampling explained less than 1% of the variance in inflammatory markers, similar to other factors including parity, tobacco use and fetal sex. Pre-pregnancy BMI and the polygenic score for CRP explained more than 9.6% and 14.1% of variance in HS-CRP levels, respectively. Together, our findings suggest an individual immune marker setpoint that is driven mostly by within-individual effects, including genetic predisposition and a metabolic component.

Our results imply that selected cytokines (IL-1β, IL-6, IL-17A, IL-23, IFN-*γ*) share similar regulatory mechanisms, while HS-CRP is likely differentially regulated. Across the four cohorts, cytokines showed moderate to strong correlation with each other, but not with HS-CRP. These findings are in line with previous studies reporting a low correlation between HS-CRP and various inflammatory markers in early pregnancy (n=110-1,274) (21,22,32) and in a non-pregnant healthy female population (33). Additional evidence for differential regulation follows from distinct patterns of cytokines and HS-CRP, as early pregnancy confidence intervals did not overlap. Specifically, cytokines showed a gradual decrease throughout early pregnancy in the current study, in line with previous studies (21–23). Cytokine levels increased towards late pregnancy in Replication Cohort I, in line with upregulation of inflammatory pathways around parturition (34). HS-CRP deviated from the cytokine pattern and showed a gradual increase followed by a decrease throughout early gestation, in line with previous studies which also showed a distinct pattern for HS-CRP and a decrease around ∼17 weeks (22) and ∼24 weeks (21). Lastly, results revealed varying drivers of cytokines and HS-CRP as discussed below. Together, our findings corroborate that HS-CRP and cytokines reflect distinct processes and may not be used interchangeably, providing important considerations for future studies in terms of immune marker selection.

Precisely timed immune adaptations have been proposed as important regulating mechanisms in successful pregnancy (23). These immune adaptations are considered to reflect trimester-specific general anti- and pro-inflammatory states (21,35). While our results showed gestational age-dependent inflammatory marker trajectories in two cohorts, we found no association with gestational age in Replication Cohort II. Across cohorts, less than 1% of the variance in inflammatory markers could be attributed to gestational age at sampling. Interestingly, when comparing pre-pregnancy and pregnancy immune markers, cytokines were also not significantly associated with gestational age. These findings suggest that pregnancy in itself may not lead to drastic cytokine changes. Additionally, distinct behavior of HS-CRP and pro-inflammatory cytokines implies a complex interaction that is poorly captured by peripheral immune markers. These findings emphasize an important gap in the extent to which preclinical findings can be translated to human pregnancies. In human cohorts, measuring systemic immune function is limited to the evaluation of clinical symptoms, such as fever or infections, or through peripheral blood sampling. How serum inflammatory marker levels relate to other immune system components and whether they reflect a one-to-one association with inflammation is still unknown. Animal studies allow for the thorough investigation of a multitude of maternal immune system components, including, but not limited to, blood biomarkers, placental function and maternal-fetal crosstalk, at different timepoints throughout gestation. To bridge findings of animal studies to human cohorts, it is essential to expand our knowledge of the function of different compartments of the immune system, and how this relates to immune biomarkers that can be measured in human pregnancies. Future preclinical studies should aim to unravel the link between blood biomarkers and underlying immune processes to better understand what these markers capture in human pregnancy.

Inflammatory markers during pregnancy were found to be driven mainly by within-individual factors, as suggested by the high intra-individual correlation between repeated measures across cohorts. Rather than a pregnancy-specific phenomenon, this has been demonstrated in non-pregnant healthy and clinical populations as confirmed by longitudinal studies (n=10-250) indicating high intra-individual immune marker reproducibility in repeated samples ranging from 7 days to 25 years apart (36–39). We identified multiple pre-pregnancy and pregnancy specific drivers of peripheral HS-CRP levels, yet together these factors explained only 12% of variance in HS-CRP levels, the majority of which was captured by pre-pregnancy BMI (9.6%). The association with parity was not reported previously (21). Our finding that female fetal sex, maternal age and tobacco use were not associated with HS-CRP is in line with previous findings (21). The association between higher BMI and increased HS-CRP was replicated in two cohorts and confirms previous reports of an association between BMI and HS-CRP (21,24,40) but less so with other cytokines (41). Given that a prior study drew from a homogeneous population of normal singleton pregnancies excluding cases of immune disorders and extreme BMI (<18 or >40), the slightly higher impact of BMI on HS-CRP observed here is likely due to the use of a population-based cohort (21). Our findings are in line with non-pregnancy studies that showed a strong correlation between BMI and HS-CRP, but not IL-6 (42,43). Several reviews have put forward CRP as a predictor of metabolic syndromes, non-alcoholic fatty liver disease and obesity, independent of inflammatory disease (40,44). While it has been suggested that chronic subclinical inflammation is a consequence of obesity (45), other studies have showed that low-grade inflammation itself may cause insulin resistance which consequently also leads to obesity and metabolic disorders such as diabetes type 2 (46). The robust association between CRP and pre-pregnancy BMI in the current pregnant population supports the interaction between inflammatory and metabolic mechanisms. We additionally identified several pre-pregnancy and pregnancy specific drivers of the maternal cytokine index – yet the total variance in cytokines explained by these factors was only 1.5%. The significant impact of maternal age, BMI and tobacco use was not shown previously, possibly due to the population-based design of the current study compared to prior reports in a healthy pregnant population (21).

Evident triggers of maternal immune activation, including bacterial or viral infections, have been associated with an upregulation of pro-inflammatory markers. Here, we investigated whether additional determinants exist that may affect the individual immune marker setpoint. Our findings revealed that infection score in early pregnancy and season of first trimester are correlated most strongly with HS-CRP and the maternal cytokine index between timepoints, respectively. Season of birth has been linked to differential maternal cytokine levels (47), as well as newborn cytokine production (48,49), possibly due to seasonality of viral exposures, vitamin D and allergen exposure (50). Our results imply that inflammatory insults and season of pregnancy may affect changes in HS-CRP, whereas HS-CRP trajectories are a reflection of genetic and metabolic components. It should be noted that effect sizes were small in comparison to the more robust drivers such as BMI and the polygenic score of CRP. Together, our findings imply that the use of cross-sectional studies/single-timepoint assessments is sufficient to identify individual baseline levels of inflammatory markers. Repeated measures of markers at different timepoints may be useful in particular for HS-CRP, considering the observed change in early pregnancy in comparison to cytokines, and to identify infection.

Our findings put into perspective the role of previously identified inflammation drivers and suggest that additional intra-individual characteristics exist which impact maternal immune marker levels, including genetic mechanisms. We therefore analyzed the polygenic score of CRP, calculated based on a recent genome-wide association study (GWAS) of CRP (28), as a predictor of HS-CRP. Our analysis revealed that the polygenic score of CRP explained a remarkable 14.1% of the variance in serum HS-CRP in the current cohort, in comparison to 16.3% of the variance in the original GWAS loci (28). In line, we showed that preconception cytokines are strongly correlated with pregnancy cytokine levels, further supporting the notion of an individual setpoint. Together, our findings are suggestive of an individual inflammatory marker setpoint which is, in part, genetically driven. Interestingly, while the release of pro-inflammatory markers is considered an orchestrated event, and CRP production is stimulated by pro-inflammatory cytokines, the polygenic score of HS-CRP revealed no association with the maternal cytokine index. Additionally, we showed that pregnancy does not lead to major changes in cytokines compared to pre-pregnancy levels, while HS-CRP is upregulated during pregnancy compared to preconception. Our findings provide further evidence that these inflammatory markers reflect distinct processes.

Strengths of the current study include the combined sample size of more than 12,000 participants in a discovery cohort, two replication cohorts, and a pre-pregnancy cohort, each including the general obstetric population, hence providing generalizable results. In addition, inflammatory marker measurement was performed by the same company using identical assays in all cohorts, minimizing technical and qualitative variance. Additionally, inflammatory markers were measured preconceptionally, providing the unique opportunity to assess pregnancy-related inflammatory marker changes. Moreover, cytokines were assessed individually as well as combined into a maternal cytokine index, to capture potential underlying mechanisms. This study is restricted by the limited knowledge of the timing of self-reported infections, which, due to the timing of questionnaires during gestation could have occurred at any point between 0-12 weeks prior to biomarker assessment. In addition to the PGS of CRP, no other PGSes of included cytokines could be used due to lack of well-powered GWASes that show high variance explained by the PGS. CRP was measured using different methodologies across the cohorts, possibly contributing to differences in median values. Lastly, as not all cytokines were measured in the replication cohorts and to maximize comparability, we assessed individual cytokines rather than a maternal cytokine index in the replication cohorts. These showed similar patterns compared to the individual cytokines in the discovery cohort. As well, varying study designs limited the option to meta-analyze the four cohorts. Yet, the application of multiple replication cohorts allowed external validation of our findings.

In conclusion, the current study mapped temporal dynamics and drivers of maternal inflammatory markers preconception and during pregnancy in the largest population to date. Our findings corroborate differential regulatory mechanisms of HS-CRP and cytokines, based on their low correlations, different trajectories, and distinct drivers throughout gestation. Additionally, our findings suggest the presence of an individual immune marker setpoint that is driven mostly by the individual, including a genetic predisposition of CRP levels and a metabolic component, rather than the pregnancy itself. Other previously identified drivers, including tobacco use and parity, explained less than 1% of inflammatory marker variance. The strong intra-individual correlation preconception and during pregnancy emphasizes the importance of using the individual as their own control when characterizing immunological profiles over time in order to discriminate between baseline inflammatory marker levels and inflammatory triggers. Taken together, further research is warranted to translate the robustness of the use of immune biomarkers in human pregnancy cohorts.

## Acknowledgements

We gratefully acknowledge the tremendous contribution of all participants of the Generation R Study, Generation C Study, Brabant Study and the Generation R *Next* Study pregnancy cohorts. We would like to thank the medical staff, biobank, and entire teams for their incredible efforts, including the recruitment, data collection and data analysis of the Generation R and Generation R *Next* cohort at the Erasmus Medical Center in Rotterdam, the Generation C cohort at the Icahn School of Medicine at Mount Sinai in New York City, and the Brabant Study cohort at Tilburg University in Tilburg. The Generation R infrastructure is located in the Erasmus MC, University Medical Center Rotterdam and functions in close collaboration with the School of Law and Faculty of Social Sciences of the Erasmus University Rotterdam, the Municipal Health Service Rotterdam area, Rotterdam, the Rotterdam Homecare Foundation, Rotterdam and the Stichting Trombosedienst and Artsenlaboratorium Rijnmond (STAR-MDC), Rotterdam. The general design of Generation R and Generation R *Next* is made possible by financial support from the Erasmus MC, Erasmus University Rotterdam, the Netherlands Organization for Health Research and Development and the Ministry of Health, Welfare and Sport.

## Materials & methods

### Study design and participants

#### Discovery cohort

The current study is embedded in the Generation R Study, a large-scale population-based prospective pregnancy cohort from early fetal life onward conducted in Rotterdam, the Netherlands. Pregnant women were recruited between April 2002 and January 2006 (51,52). The enrollment procedure has been described elsewhere (53). The Generation R study was approved by the Medical Ethical Committee (MEC 198.782/2001/31) of the Erasmus Medical Center, Rotterdam, the Netherlands. All participants provided written informed consent. Of 9,778 participants enrolled during pregnancy, 72.3% were enrolled prior to 18 weeks gestation (n=7,069). Participants were excluded from the current analysis if no cytokine measurement at any point during gestation was available (n=1,644, 16.8%), if the assay showed low bead count or if there was insufficient sample volume available (n=53, 0.5%). A total of 8,082 participants were included in the current study. The study design is shown in **Figure 1A**.

#### Replication Cohort I

The first replication cohort is the Generation C Study conducted at the Mount Sinai Hospital in New York City, USA (54). In short, pregnant individuals who received obstetrical care between April 2020 and February 2022 in the Mount Sinai Health System were eligible for participation. The study was approved by the Icahn School of Medicine at Mount Sinai Institutional Review Board (IRB-20-03352), reviewed by the US Centers for Disease Control and Prevention (CDC) and consistent with applicable federal law and CDC policy. All participants provided informed consent. In total, 3,157 participants were included in the Generation C Study. For the current study, participants were included if at least one cytokine measurement at any point during gestation was available (n=2,535, 80.3%).

#### Replication Cohort II

The second replication cohort was the Brabant Study conducted at various midwife practices in the Netherlands (55). In short, pregnant individuals were recruited at 8-10 weeks gestation among midwifery practices in the South-East of the Netherlands between June 2018 and January 2023. All participants provided informed consent. The Brabant study was approved by the Medical Ethical Committee of the Máxima Medical Center Veldhoven (protocol number NL64091.015.17). In total, 2863 participants were included in the Brabant Study. High-sensitivity cytokine assays were performed in a subset of participants. For the current study, participants were included if at least one cytokine measurement at any point during gestation was available (n=587, 20.5%).

#### Pre-pregnancy Cohort

The Generation R *Next* study cohort is a population-based prospective cohort study from preconception onwards, conducted in Rotterdam, the Netherlands. Recruitment started in August 2017 and is currently ongoing. Participants of reproductive age planning to have children are recruited at general practitioners. Pregnant women are invited to the research center in the first and third trimester. The Generation R *Next* study was approved by the Medical Ethical Committee (MEC-2016-589, December 2016) of the Erasmus Medical Center, Rotterdam, the Netherlands. All participants provided written informed consent. A total of 1,270 participants were included in the current study with at least one cytokine measurement available preconception or in the first trimester.

### Sample collection

#### Discovery cohort

In the Generation R Study, maternal venous blood samples were collected as part of routine prenatal visits at the midwife practice with the first blood sample collected at median 13.2 weeks (95% range: 9.6-17.6 weeks, n=6,072) and the second blood sample at median 20.4 weeks (95% range: 18.5-23.3 weeks, n=7,395), with an overall median of 19.2 weeks (95% range: 10.6 – 23.1 weeks, n=13,467). Blood was collected through ante-cubital venous puncture, temporarily stored at room temperature and transported to the regional laboratory (Star-MDC, Rotterdam, the Netherlands) within three hours of collection (52). Blood samples were centrifuged for 10 minutes, and the serum was aliquoted into 250ul into polypropylene Micronic tubes, and immediately stored at -80°C. Freeze-thaw cycles were avoided. Samples were bar coded with a unique laboratory number.

#### Replication Cohort I

In the Generation C Study (Replication Cohort I), blood specimens were obtained as part of routine blood draws during prenatal visits or at admission to the labor and delivery unit. Blood samples were collected at median 27.7 weeks (95% range: 6.9-40.6 weeks, n=3,319). Blood was collected through ante-cubital venous puncture, temporarily stored at room temperature and processed within 4 hours (median = 1.6 hours; IQR = 1.7 hours). Blood samples were centrifuged for 10 minutes, aliquoted into 500 µl vials and stored at -80°C until further analysis. Freeze-thaw cycles were avoided. Samples were bar coded with a unique laboratory number.

#### Replication Cohort II

In the Brabant Study (Replication Cohort II), blood sample collection was performed by the regional organization responsible for blood collection for primary and secondary care laboratories (De Bloedafname). Blood samples were collected at three timepoints during gestation, namely at 12, 20 and 28 weeks, with an overall median of 20.2 weeks (95% range: 12-29.5 weeks, n=1,170). Blood samples were processed and stored by the laboratory of clinical chemistry and hematology of the Máxima Medical Centre Veldhoven. Freeze-thaw cycles were avoided. Samples were bar coded with a unique laboratory number.

#### Pre-pregnancy Cohort

In the Generation R *Next* Study (Pre-pregnancy Cohort), blood sample collection was performed during an appointment at the research center prior to pregnancy at median 11.4 weeks prior to conception (95% range: 74.5-3.6 weeks prior to conception, n=676), and two blood samples during pregnancy in the first trimester at median 8.4 weeks gestation (95% range: 6.4-12.9 weeks, n=1,103) and in the third trimester at median 30.3 weeks gestation (95% range: 29-33.6 weeks, n=1,067). Blood collection and processing procedures were very similar to the Discovery Cohort. For the current analyses, only preconception and first trimester samples were included.

### Inflammatory marker measurement

#### Discovery cohort

##### High-sensitivity C-reactive Protein (HS-CRP)

A high sensitivity assay was used to measure HS-CRP to increase the sensitivity in the low range with improved precision at low CRP concentrations (56). HS-CRP assays were performed by the Department of Clinical Chemistry of the Erasmus MC, using an immunoturbidimetric assay on the Cobas 6000 analyzer (Roche Diagnostics). The lowest level of detection was 0.2 mg/L. The early and mid-pregnancy samples were run in two separate batches (2006 and 2022), using different assay kits and lot numbers. HS-CRP measurement of the first batch of mid-pregnancy blood samples has been described elsewhere (57).

##### Pilot: cytokine selection

In our search for factors that impact the maternal immune system during pregnancy, we first set out to construct a measure of maternal immune activation. Given that immune markers interact and are involved in specific pregnancy processes, it is of interest to quantify a panel of robust cytokines that have the potential to capture both low-grade and acute systemic inflammation. Quantification of 14 cytokines (GM-CSF, IL-1β, IL-2, IL-4, IL-5, IL-6, IL-8, IL-10, IL-12p70, IL-13, IL-17A, IL-23, IFN-*γ* and TNF-*α*) as part of a pilot (n=100) revealed excellent performance with no values below the detection limit. Several outliers were excluded (**Supplementary Figure 1B**). Clustering analyses showed correlations between cytokines including IL-1β, IL-2, IL-4, IL-6 IL-13, IL-17A and IFN-*γ*, while TNF-*α* and IL-8 did not correlate with other cytokines (**Supplementary Figure 1A**). Cytokines were differently distributed, with IL-6 showing values in the low range and IL-23 showing values in a higher range (**Supplementary Figure 1C**). This data-driven approach led to the selection of cytokines that correlate highly with each other and capture both markers in the low and in the higher range. Next, based on literature suggesting a role in pregnancy processes in particular for IL-1β, IL-6, IL-17A, and IFN-*γ* (18–20), a final selection was made. Inflammatory cytokines IL-1β, IL-6, IL-17A, IL-23 and IFN-*γ* were measured in the entire cohort to establish inflammatory marker patterns and a marker of immune activation in early and mid-pregnancy (**Supplementary Figure 1C**).

##### Processing and quality assurance of cytokine data

Cytokine analyses were performed by Eve Technologies Corp., Calgary, Canada. A pilot study of 100 serum samples was performed using the Human High Sensitivity 14-Plex Discovery Assay which simultaneously quantified levels of 14 cytokines (granulocyte-macrophage (GM-) colony-stimulating factor (CSF), interferon (IFN)-*γ*, interleukin (IL)-1β, IL-2, IL-4, IL-5, IL-6, IL-8, IL-10, IL-12p70, IL-13, IL-17A, IL-23, and tumor necrosis factor (TNF)-*α* (pg/mL). Pilot results showed 100% detection within the standard curve, indicating high sensitivity and high assay performance. IL-1β, IL-6, IL-17A, IL-23, and IFN-*γ* were measured in the full Generation R cohort using the Human High Sensitivity T-Helper Cells Custom 5-plex assay from Millipore (Millipore, St. Charles, MO, USA) on the Luminex™ 100 system (Luminex, Austin, TX, USA). Serum samples were analyzed across 145 plates in 37 batches. Maternal and fetal characteristics including maternal age, BMI, fetal sex, parity, and birthweight were randomly distributed across plates (**Supplementary Figure 1E**). Several measures were taken to harbor the quality and limit variance based on laboratory conditions, namely: i) quality control (QC) samples were included in each assay session. All QC samples used in the study were from the same lot number, and reconstituted and aliquoted per kit protocol at the start of the study; ii) all analyses were run on the same kit and lot specific reagents (Kit Lot number 3891089; Detection Antibodies Lot number 3692805); iii) to minimize potential intra-assay variables, all analyses were run on the same analyzer by the same technician, and iv) ancillary fluidic components including calibration material and sheath fluid were from the same lot number for the entire study. We found no effect of cytokine batch (**Supplementary Figure 1D**), however, to avoid noise, batch was added as a covariate in the main analyses. The observed analyte concentrations are generated based on cubic spline regression analyses using the expected concentration values of the standard, as defined in the manufacturer’s instructions for use, and the Fluorescence Intensity (FI), as suggested by Breen et al. (58). Cytokine concentrations were estimated from the FI using specific calibration curves for each analyte, constructed from standard samples in the reference lot. Less than 0.5% of the samples fell outside of the standard curve on the low end based on the cubic spline curve for that specific analyte. These values were substituted with the lowest measured value for that specific analyte (59). Immunoassay performance is shown in **Supplementary Table 10**. Cytokines IL-1β, IL-6, IL-17A, IL-23, IFN-*γ* and HS-CRP were log2 transformed, to satisfy the normality assumption for downstream analyses.

##### Potential HAAA interference

Circulating human antibodies that are reactive with animal antibodies (human anti-animal antibodies, or HAAA), may pose a source of immunoassay interference. HAAA are high affinity, specific polyclonal antibodies produced by the human immune system against a specific animal immunogen (heterophile antigen), or due to non-iatrogenic causes such as pharmaceutical agents or vaccines (60). The most encountered HAAA are human anti-mouse antibodies (HAMA), considered to be present in about 5-10% of the population, with a potential increase in an inflammatory population as antibody-based therapeutics might be more frequent. In samples with HAMA, the anti-mouse antibodies can bind and cross-link capture antibodies to detection antibodies in the absence of the analyte, resulting in artificially high fluorescence values in immunoassays based on mouse-antibodies, such as the assay that we used (61). In the Milliplex assay, potential HAMA interference is indicated in samples with a median fluorescent intensity value above 500 across the five cytokines. Characteristics of inflammatory markers with and without HAAA samples are shown in **Supplementary Table 11**. However, as we are interested in states of immune activation, where increased cytokine levels are to be expected, we further investigated the behavior of samples with possible HAMA interference in a sensitivity analysis.

##### Replication Cohorts and Pre-pregnancy Cohort

In both replication cohorts and the pre-pregnancy cohort, IL-1β, IL-6 and IL-17A were assessed using the High Sensitivity T-cell Discovery Array 3-Plex (Millipore, St. Charles, MO, USA) at Eve Technologies using the Bio-Plex™ 200 system (Bio-Rad Laboratories, Inc., Hercules, CA, USA). For the pre-pregnancy cohort, IL-23 and IFN-*y* were also measured as part of this assay. For Replication Cohort I, HS-CRP was also analyzed as part of this assay. For Replication Cohort II, CRP was measured by the laboratory of clinical chemistry and hematology of the Máxima Medical Centre Veldhoven. Similar processing and quality assurance checks were performed as the discovery cohort. For the pre-pregnancy cohort, HS-CRP was measured by the Department of Clinical Chemistry of the Erasmus MC, similar to the Discovery Cohort.

### Pre-pregnancy Characteristics and Pregnancy Circumstances

#### Data collected at enrollment (pre-pregnancy characteristics)

Questionnaires at enrollment were used to assess maternal demographic variables, including maternal age at conception, pre-pregnancy body mass index (BMI) (kg/m^2^), education level (no education, primary education, secondary education), national background (Dutch, Non-Dutch), parity (nulliparous, multiparous), immune-related disease (HIV, eczema, systemic lupus erythematosus (SLE), intestinal disorder, pre-gestation diabetes, multiple sclerosis, rheumatism) and household income (low: <€2,220/month or high: >€2200/month, based on the average household income in Rotterdam in 2005). In addition, prenatal maternal psychopathology was assessed at enrollment using the Brief Symptom Inventory (62). A Global Severity Index (GSI) was calculated indicating prenatal maternal psychopathology with higher scores meaning more problems and referred to as ‘Maternal Psychopathology’ throughout the manuscript. These pre-existing maternal characteristics are referred to as pre-pregnancy characteristics.

#### Data collected during pregnancy (pregnancy circumstances)

Data on maternal tobacco use (never smoked, tobacco use pre-pregnancy, tobacco use during pregnancy), alcohol consumption (never drank, alcohol use pre-pregnancy, occasional alcohol use during pregnancy, frequent alcohol use during pregnancy) and substance use during pregnancy (yes/no), was obtained through questionnaires collected at three times during pregnancy. Questionnaire data obtained prior to or at the time of blood sample collection was included. Information on fetal sex was obtained from medical records. Gestational age of the sample was analyzed as a continuous variable, and further categorized into season following European references (spring, summer, fall, winter). These pregnancy-specific characteristics are referred to as pregnancy circumstances.

#### Infection score

Questionnaire data collected at three times during pregnancy was used to create a self-rated continuous infection score, as described elsewhere (63). An infection score of 0 indicates no infection was reported. An infection score of 1 and higher indicates the participant reported any one of the following infections at least once: upper respiratory infections (pharyngitis, rhinitis, sinusitis, ear infection), lower respiratory infections (pneumonia, bronchitis), gastrointestinal infections (diarrhea, enteritis), cystitis/pyelitis, dermatitis (boils, erysipelas), eye infections, herpes zoster, flu, sexually transmitted diseases (STD), and a period of fever (>38°C/100.4°F) in the 2-3 months prior to blood draw.

#### Replication Cohort I

In Replication Cohort I, information on maternal demographic variables was obtained through questionnaires collected at enrollment. These included maternal age at conception, pre-pregnancy body mass index (BMI) (kg/m^2^), national background (white, non-White), parity (nulliparous, multiparous) and insurance status (private/self-pay, public). These pre-existing maternal characteristics are referred to as pre-pregnancy characteristics. Information on fetal sex, twin pregnancy, birthweight and gestational age at birth was obtained from medical records. These pregnancy-specific characteristics are referred to as pregnancy circumstances.

#### Replication Cohort II

In Replication Cohort II, information on maternal demographic variables was obtained through questionnaires collected at each visit at 12, 20, and 28 weeks. These included maternal age at conception, pre-pregnancy body mass index (BMI) (kg/m^2^), national background (Dutch, non-Dutch), parity (nulliparous, multiparous), education level (no education, primary education, secondary education), immune-related disease (HIV, eczema, systemic lupus erythematosus (SLE), intestinal disorder, pre-gestation diabetes, multiple sclerosis, rheumatism) and employment status (employed, unemployed). These pre-existing maternal characteristics are referred to as pre-pregnancy characteristics. Information on fetal sex, birthweight and gestational age at birth was obtained from medical records. These pregnancy-specific characteristics are referred to as pregnancy circumstances.

### Computation of Polygenic Score (PGS) of CRP

#### Discovery cohort Genotype data

Maternal samples were used for DNA extraction, as detailed elsewhere (52). Parents in Generation R were genotyped in two batches using the Illumina Global Screening Multi-Disease Array (GSA-MD) v2 (1,530 mothers in 2019/2020) and GSA-MD v3 (10,491 mothers and fathers in 2022) platforms. Both batches underwent extended quality control checks. Genotype call rate was checked in two rounds, the initial with a threshold of 95% and then with 97.5%. Single nucleotide variants (SNVs) that failed the one-sided HWE test (p≤1*10-5) were removed, as well as samples with evidence of excess heterozygosity, gender mismatch, unexpected genetic duplicates and with unexpected familial bonds. After the extended quality control checks, the batches were merged, leaving 11,742 parents (including 7,256 mothers) and 660,868 SNVs. After merging the two batches, the unmapped single nucleotide polymorphisms (SNPs) were imputed to a reference panel (1000 Genomes Project - phase3 version 5, build 38) by integrating SHAPEIT and Minimac4 programs in an in-house *Odyssey* pipeline. The imputation quality threshold was set to 0.8 and the MAF threshold was set to 0.01.

#### CRP polygenic score (PGS)

A recent large-scale gene-wide association study (GWAS) with 575,531 participants was used to construct a PGS of CRP based on imputed genotypic data (28). The summary statistics were obtained from the GWAS study catalog (ID: GCST90029070). The PGS was calculated for participants of any ancestry (n=5,938) and for participants of European descent (n=4,238) separately. LDpred2, the latest version of LDpred that offers better and more robust predictions, was used to calculate the PGS (64,65). LDpred2 is implemented within the R-package *bigsnpr* (66) and derives PGSes based on summary statistics and Linkage Disequilibrium (LD) information from an external reference panel (65). The variants in the summary statistics were matched to the ones in the LD reference and the maternal genotypes. The final dataset included 1,044,979 SNPs. We used a genome-wide BED file with the maternal genotypes, HapMap3 variants with individual LD matrix in blocks and LDpred2-auto (one variant of LDpred2) to automatically estimate p (the proportion of causal variants) and *h^2^*(the SNP heritability) from the summary statistics. After estimating *h^2^*, LDpred2-auto was run with 30 iterations. A sequence of 30 values from 10−4 to 0.2 equally spaced on a log scale were used as initial values for p. A shrinkage coefficient of 0.95 was used and effects sizes were forced to go through 0 first before changing sign in consecutive iterations to prevent instability of the Gibbs sampler as suggested by Privé (67). To get the final effects, only chains that passed the quality control filtering were used. The calculated score was then standardized to z-scores (mean 0, 1 SD) and residualized on the first 10 principal components. The R^2^ change between the null model and the model including the PGS of CRP is shown in **Supplementary Table 12**. A higher PGS of CRP indicates a higher genetic risk for elevated CRP levels. Said et al. (2022) showed that the independent variants within the UK Biobank GWAS loci explained 16.3% of the variance in CRP levels. Prior studies have shown that the PGS of CRP is associated with cardiovascular outcomes and performs well in multiple cohorts (68,69).

#### Replication Cohorts

No genotyping data was available.

### Computation of maternal cytokine index

#### Discovery cohort

Cytokines are part of complex networks. To account for immune marker inter-relationships, a composite score of inflammatory markers was composed for each sample: the maternal cytokine index. Collapsing the inflammatory marker data into a maternal cytokine index allows to capture the underlying structure of the cytokine data and provide a comprehensive summary of cytokine patterns. Principal component analysis (PCA) was used as a means of dimensionality reduction using the ‘prcomp’ function from the ‘stats’ package in R. The principal components represent linear combinations of the cytokine data. A higher loading of a cytokine on a principal component indicates increased contribution of that specific cytokine to the overall variability. In addition, PCA addresses multicollinearity by orthogonalizing the cytokines into uncorrelated principal components. This improves the stability and reliability of subsequent analyses. PCA revealed two distinct inflammatory marker patterns. On the one hand, cytokines IL-1β, IL-6, IL-17A, IL-23, and IFN-*γ* loaded highly on the first principal component (PC) (84%, 66%, 89%, 77%, 89%, respectively), accounting for 55% of the variance. In turn, CRP loaded highly on the second PC (98%). Additionally, IL-6, but not other immune markers, loaded on the third PC (68%). Adding the third PC did not substantially improve model fit and we did not consider the third PC to reflect a profile of biological relevance. An elbow plot suggested that two PCs should be taken into account. Hence, only the first two PCs were considered. Cytokines IL-1β, IL-6, IL-17A, IL-23, and IFN-*γ* were used to create the maternal cytokine index, defined as the eigenvalue, or singular value decomposition, aggregated across the cytokines, equivalent to the first principal component. Due to its high loading on the second PC, HS-CRP was not included in the maternal cytokine index and was assessed separately. The maternal cytokine index was included as a continuous score in analyses.

#### Replication Cohorts

Not all cytokines of the discovery cohort panel were measured in the replication cohorts and no maternal cytokine index was created in the replication cohorts. Individual cytokines were assessed to maximize comparability to the discovery cohort.

### Statistical analysis

Analyses were performed using the R Statistical Software (version 4.1.2) (R Core Team, 2020). Visualizations were made with the ggplot2 package (70). Visual representation of study design was created using Biorender.

#### Demographics

Demographic characteristics were presented as means with standard deviation (SD) for normally distributed continuous data, medians with inter-quartile range (IQR) for non-normally distributed continuous data and as numbers (percentages) for categorical data.

#### Outlier analysis

Outlier samples for inflammatory markers were identified using unsupervised hierarchical clustering, based on Pearson coefficient and average distance metric, and principal component analysis (PCA). Samples more than three SD from the grand mean of the first PC were considered potential outlier samples. Considering that these samples might reflect levels of acute or chronic inflammation, they were included in the main analysis and excluded in a sensitivity analysis.

#### Missing data and imputation

In the Discovery Cohort, covariates with missing data included parity (1%), national background (5%), education (8%), alcohol use (10%), tobacco use (12%) and substance use (12%), pre-pregnancy BMI (18%), maternal psychopathology (22%) and household income (22%). In Replication Cohort I, missing data included insurance status (0.2%) and fetal sex (10.4%). In Replication Cohort II, missing data included education level (1.9%), parity (1.9%), employment status (2.0), tobacco use (2.2%), alcohol use (2.2%) and fetal sex (10.7%). Missing covariate data were imputed using the Mice package in R (100 iterations, 30 datasets) and analyses were conducted across the pooled datasets following Rubin’s rules.

#### Aim 1: mapping dynamics throughout gestation

Inflammatory marker levels were compared between timepoints using an unpaired t-test. Correlations among inflammatory markers were assessed using Pearson’s r correlation coefficient across all samples, as well as at timepoint 1 and 2 separately. In addition, the correlation between inflammatory markers and the maternal cytokine index was assessed across all samples. Correlations of inflammatory markers between timepoint 1 and 2 were assessed pairwise among participants with repeated measures. Inflammatory marker trajectories were modeled with (non-)linear generalized additive models (GAM) (formula = y ∼ s[x, bs = “cs”]) using the ‘ggplot’ package in R. Inflammatory markers were centered and scaled by subtracting the column means from each sample and dividing by their standard deviation using the ‘scale’ function in R. Next, variance partition analysis was performed to assess potential drivers of the variation in inflammatory marker expression using the ‘variancePartition’ package in R (71). Categorical predictors were modeled as random intercepts and continuous predictors were modeled as fixed effects. Collinearity between predictors was assessed with the ‘fitVarPartModel’ function.

#### Aim 2: drivers of maternal inflammatory markers

Linear mixed-effects regression models were applied to investigate the association between potential drivers (exposures) and HS-CRP and the maternal cytokine index (outcomes) in separate models. LMM were performed with log-normalized inflammatory markers as outcome variables. Gestational age at sampling and cytokine batch were included as fixed effects. Participant was included as random intercept to account for repeated measures from the same participant. Three separate models were run; model 1 includes maternal age, pre-pregnancy BMI, PGS of CRP, national background, household income, parity, maternal psychopathology, tobacco use, alcohol use, substance use, fetal sex, and season of first blood draw as fixed effects. The infection score was interpreted from a separate model as we hypothesize that BMI may be on the pathway of pregnancy infection and inflammatory markers (Model 2) (72,73). Models 1 and 2 were run in a subset of participants with complete CRP PGS for all ancestry (n=5,938 participants, n=10,157 samples). The PGS of European ancestry was interpreted from a separate model among participants with a complete PGS of European ancestry (n=4,238, n=7,381 samples) (Model 3).

The linear mixed effects regression model equations were:

- Model 1: Maternal cytokine index / HS-CRP / Individual cytokines ∼ maternal age + pre-pregnancy BMI + CRP PGS (all ancestry) + national background + household income + parity + maternal psychopathology + tobacco use + alcohol use + substance use + fetal sex + season of first blood draw + cytokine batch + gestational age of the sample + (1 l participant_i_) + ε_i._
- Model 2: Maternal cytokine index / HS-CRP / Individual cytokines ∼ maternal age + pre-pregnancy BMI + CRP PGS (all ancestry) + national background + household income + parity + maternal psychopathology + tobacco use + alcohol use + substance use + fetal sex + season of first blood draw + infection score + cytokine batch + gestational age of the sample + (1 l participant_i_) + ε_i._
- Model 3: Maternal cytokine index / HS-CRP / Individual cytokines ∼ maternal age + pre-pregnancy BMI + CRP PGS (European ancestry) + national background + household income + parity + maternal psychopathology + tobacco use + alcohol use + substance use + fetal sex + season of first blood draw + cytokine batch + gestational age of the sample + (1 l participant_i_) + ε_i._

The ‘i’ subscript indicates the participant. The R packages lme4 and lmertest were used to fit the models (74). Linear mixed-effects models were fit with the restricted maximum likelihood as estimation method. Assumptions were checked for all models. Collinearity between variables in the models was assessed based on the variance inflation factor (VIF). If the VIF>3, variables were considered to be collinear. All variables were below VIF =2. Benjamini-Hochberg correction was applied per outcome and per aim to correct for multiple testing (75). Both unadjusted and corrected p-values are shown in the tables. Statistical significance is considered if *q*<0.05 after multiple testing correction. P-values reported within the manuscript are corrected for multiple testing.

#### Delta between timepoints

The change in inflammatory marker levels between timepoints can be used to provide insight in inflammatory triggers that occurred between two timepoints. A continuous delta was computed for HS-CRP and the maternal cytokine index by subtracting the measurement of the first timepoint from the measurement of the second timepoint. Canonical correlation analysis was performed to assess the correlation between predictors and the delta HS-CRP and delta maternal cytokine index. Canonical correlation analysis was performed using the ‘variancePartition’ in R which allows the inclusion of both continuous and discrete variables in one formula. Additionally, participants were divided into three groups based on the HS-CRP delta and the maternal cytokine index delta. The groups were defined as the <5^th^, 5^th^-95^th^ and >95^th^ quantiles of the HS-CRP delta and maternal cytokine index delta, resulting in a high-low (high levels at T1 and low at T2; 5^th^ quantile of the delta), stable, and low-high group (low levels at T2 and high at T2; 95^th^ quantile of the delta). A sensitivity analysis was performed to determine the associations between predictors and HS-CRP and the maternal cytokine index among samples of participants with a high absolute HS-CRP delta (n=435 participants; n=854 samples).

#### Sensitivity analyses

Several sensitivity analyses were performed: i) excluding samples with potential HAAA interference (n=123 participants; n=163 samples), to determine the associations without possible HAAA interference; ii) excluding samples of participants with immune-related diseases (defined as having any one of the following immune-related diseases: HIV, eczema, systemic lupus erythematosus (SLE), intestinal disorder, pre-gestation diabetes, multiple sclerosis, rheumatism), to determine the associations without potential noise due to inherent immune dysregulation (n=495 participants; n=855 samples); and iii) excluding samples considered outliers based on the outlier analysis described above (n=147 participants; n=174 samples), to determine the associations without cases that may unduly influence the inflammatory marker estimates; and iv) for each cytokine as an outcome, to determine associations between predictors and individual cytokines.

---

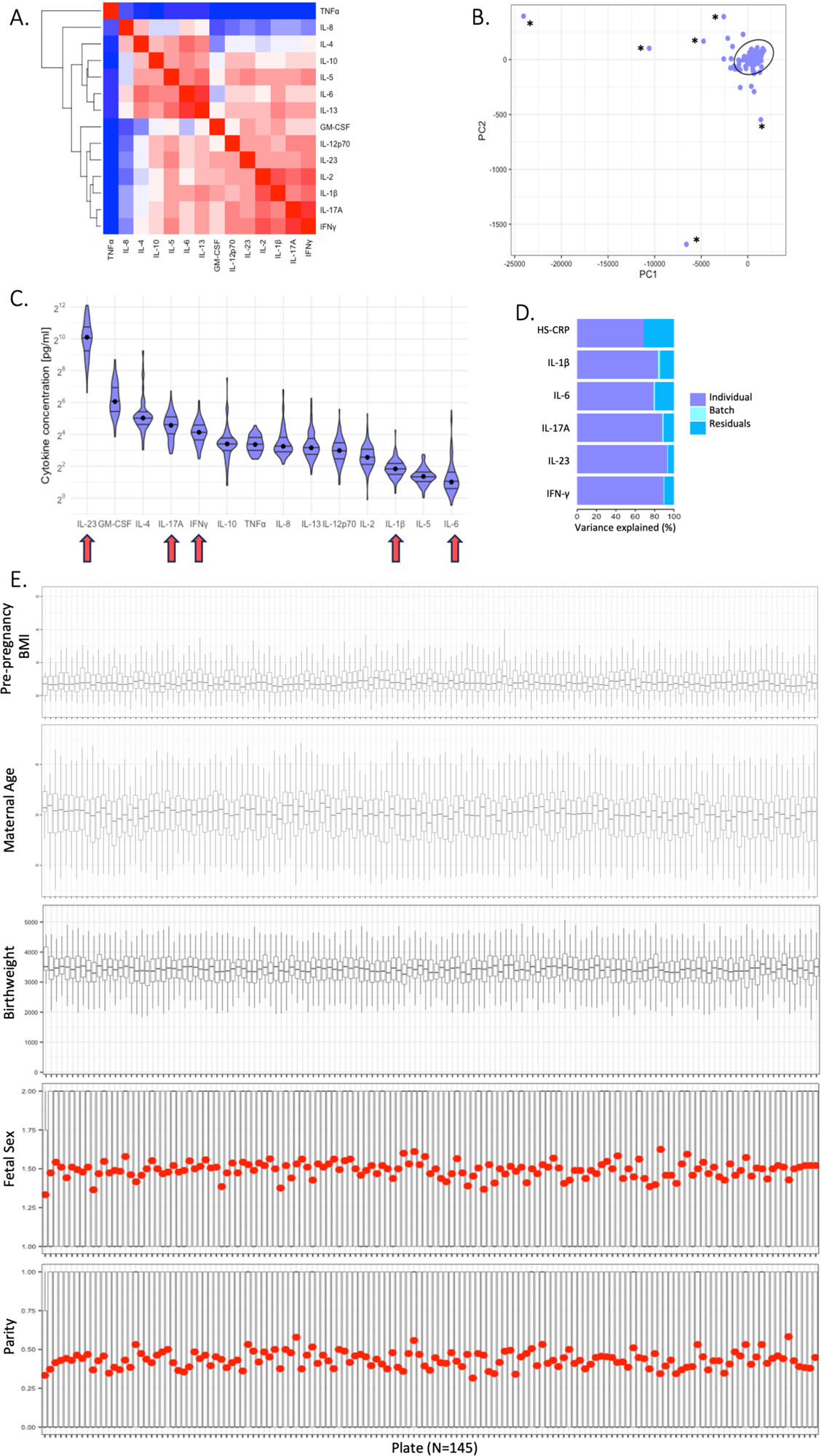

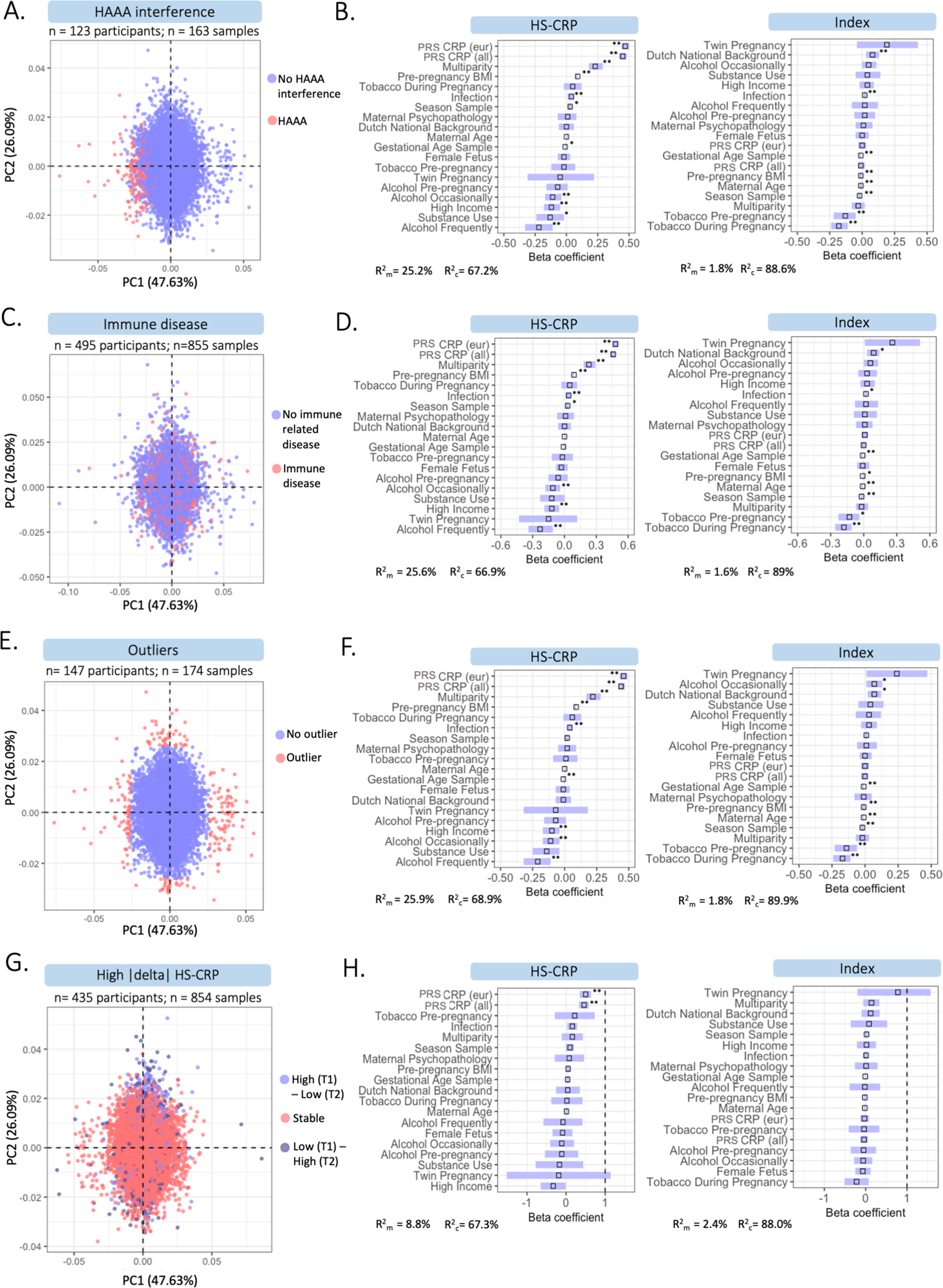

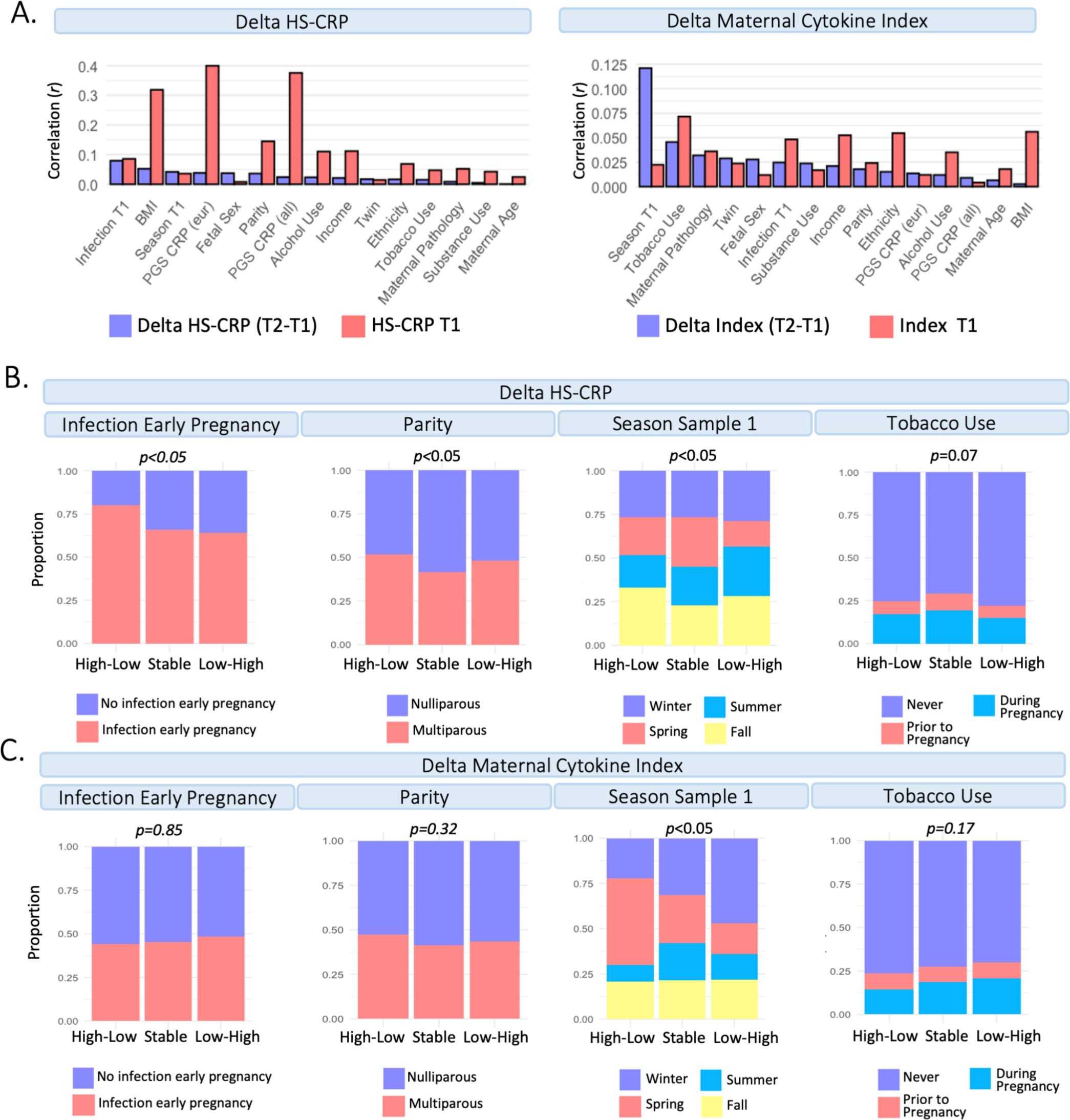

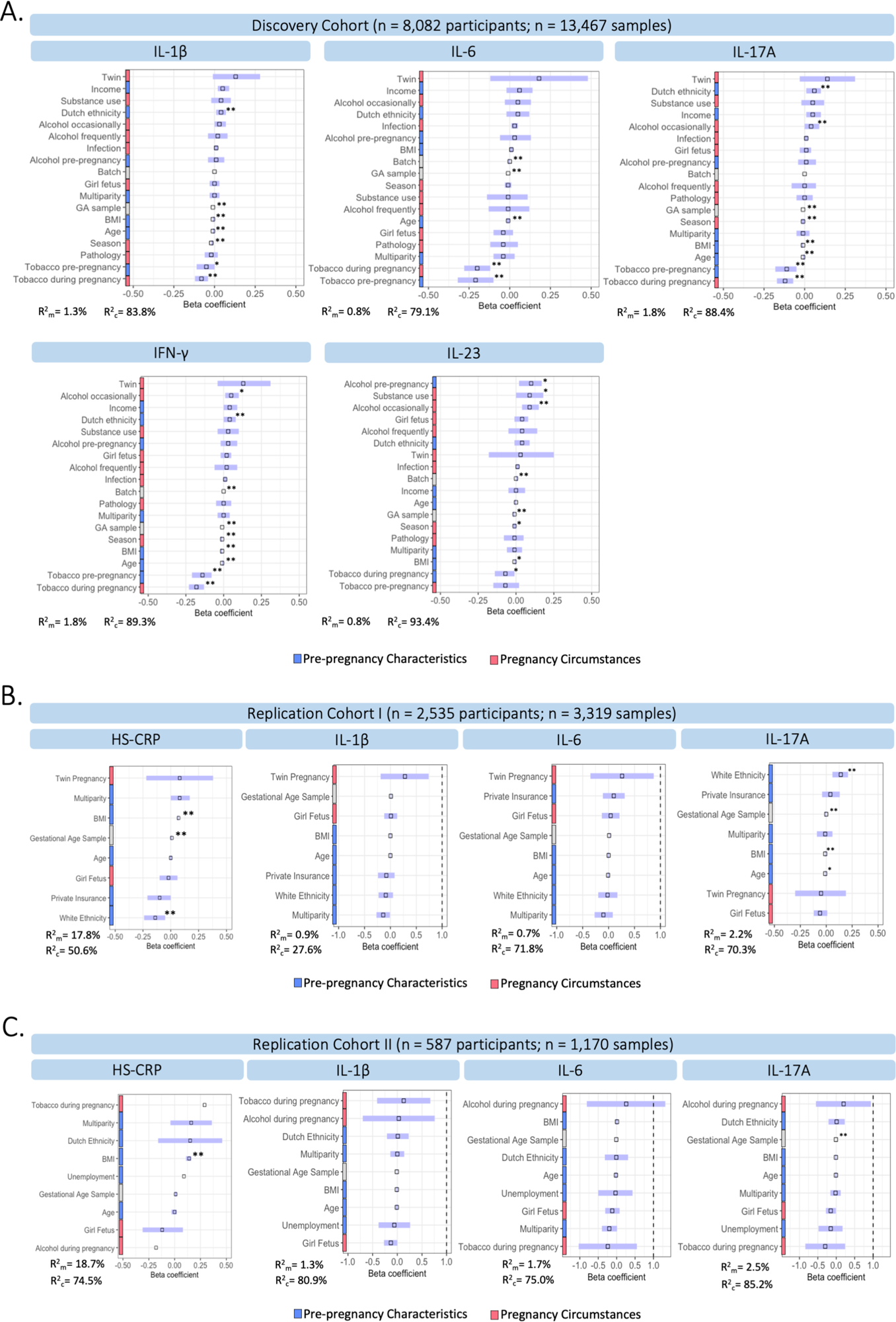

