## Supplementary Materials for "Inflammatory markers in pregnancy – surprisingly stable. Mapping trajectories and drivers in four large cohorts"

**Supplementary Figure 1. Multiplex analysis of cytokines in the Discovery Cohort.** A) Clustering of 14 cytokines measured in a multiplex assay pilot analysis (n=100). B) Principal component analysis (PCA) showed six outlier samples, indicated by an asterisk. C) The distribution of 14 cytokines in pg/ml, on a log2 y-axis. The dot indicates the median, the lines indicate the 25^th^, 50^th^ and 75^th^ percentile, respectively. Cytokines selected in the final panel are indicated with an arrow. D) Variance explained in HS-CRP and cytokines by technical variable cytokine batch. E) Several potential drivers were randomized across 145 plates prior to multiplex analysis.

**Supplementary Figure 2. Multivariable associations between potential drivers and inflammatory markers in sensitivity analyses.** A) Distribution of samples with potential human anti-animal antibodies (HAAA) interference (n=163) across Principal Component Analysis (PCA). B) Mixed effects linear regression models of potential drivers and their association with HS-CRP (left) and the maternal cytokine index (right) after excluding samples with potential human anti-animal antibodies (HAAA) interference (n=163). C) Distribution of samples of participants with auto-immune disease (n=855) across Principal Component Analysis (PCA). D) Mixed effects linear regression models of potential drivers and their association with HS-CRP (left) and the maternal cytokine index (right) after excluding samples of participants with auto-immune disease (n=855). E) Distribution of outlier samples more than 3 SD from the grand mean of PC1 and PC2 (n=174) across Principal Component Analysis (PCA). F) Mixed effects linear regression models of potential drivers and their association with HS-CRP (left) and the maternal cytokine index (right) after excluding outlier samples (n=174). G) Distribution of samples of participants with a high absolute HS-CRP delta (n=854) across Principal Component Analysis (PCA). H) Mixed effects linear regression models of potential drivers and their association with HS-CRP (left) and the maternal cytokine index (right) in a subgroup of samples of participants with a high absolute HS-CRP delta (n=854). Legend: R^2^_m_ indicates the marginal R squared of the mixed effects regression model, R^2^_c_ indicates the conditional R-squared of the mixed effects regression model after adding individual as a random intercept. Asterisks indicate significance level after multiple testing correction (*p<0.05, **p<0.01, ***p<0.001).

**Supplementary Figure 3. Change in inflammatory markers between timepoints.** A) Correlation between potential drivers and the HS-CRP delta (HS-CRP timepoint 2 - HS-CRP timepoint 1) and HS-CRP levels at timepoint 1 (left). Correlation between potential drivers and the maternal cytokine index delta (maternal cytokine index timepoint 2 – maternal cytokine index timepoint 1) and the maternal cytokine index at timepoint 1 (right). Canonical correlation coefficients (Pearson’s r) are shown. B) Participants with repeated measures were categorized based on the HS-CRP delta and divided into a high-low (high levels at T1 and low at T2; 5^th^ quantile of HS-CRP delta), stable, and low-high group (low levels at T1 and high at T2; 95^th^ quantile of HS-CRP delta). The *p*-value indicates whether the proportion differed significantly between groups. C) Participants with repeated measures were categorized based on the maternal cytokine index delta and divided into a high-low (high levels at T1 and low at T2; 5^th^ quantile of maternal cytokine index delta), stable, and low-high group (low levels at T1 and high at T2; 95^th^ quantile of maternal cytokine index delta). The *p*-value indicates whether the proportion differed significantly between groups.

**Supplementary Figure 4. Multivariable associations between potential drivers and inflammatory markers in three cohorts.** A) Mixed effects linear regression models of potential drivers and their association with IL-1β, IL-6, IL-17A, IFN-*γ* and IL-23 in the Discovery Cohort (n=13,467 samples)**.** B) Mixed effects linear regression models of potential drivers and their association with HS-CRP, IL-1β, IL-6 and IL-17A in Replication Cohort I (n=3,319 samples). C) Mixed effects linear regression models of potential drivers and their association with HS-CRP, IL-1β, IL-6 and IL-17A in Replication Cohort II (n=1,170 samples). Legend: Forest plots show beta coefficients and 95% confidence interval of selected predictors with each inflammatory marker. Pre-pregnancy characteristics are indicated in blue, pregnancy circumstances are indicated in red.

**Supplementary Table 1: Characteristics of inflammatory markers in Replication Cohort I.** Inflammatory marker levels overall, as well as per timepoint in maternal serum samples obtained from n=2,535 participants in Replication Cohort I.

| **Inflammatory marker** | **Overall (n=3,319)** | | **Timepoint 1***  **(n=1,290)** | | **Timepoint 2***  **(n=2,029)** | | **p-value**** |
| --- | --- | --- | --- | --- | --- | --- | --- |
|  | **Median (IQR)** | **Range (min-max)** | **Median (IQR)** | **Range (min-max)** | **Median (IQR)** | **Range (min-max)** |  |
| IL-1β (pg/ml) | 2.38 (3.75) | 0.02-376.34 | 2.41 (3.00) | 0.03-247.88 | 2.35 (4.39) | 0.02-376.34 | **0.025** |
| IL-6 (pg/ml) | 1.24 (2.42) | 0.01-155.46 | 1.21 (2.04) | 0.01-155.46 | 1.25 (2.77) | 0.01-82.12 | 0.066 |
| IL-17A (pg/ml) | 9.85 (6.50) | 0.80-244.18 | 10.30 (6.75) | 1.05-220.61 | 9.56 (6.27) | 0.80-244.18 | **<0.001** |
| HS-CRP (mg/L) | 16.9 (15.96) | 0.20-408.36 | 15.45 (17.88) | 0.20-204.23 | 17.58 (14.76) | 0.32-408.36 | **<0.001** |

*Timepoint according to study design: Timepoint 1 = early pregnancy, $\leq$20 weeks gestation. Timepoint 2 is late pregnancy, >20 weeks gestation.

**Paired samples t-test of normalized cytokines and HS-CRP in early and late pregnancy.

**Supplementary Table 2: Characteristics of inflammatory markers in Replication Cohort II.** Inflammatory marker levels overall, as well as per timepoint in maternal serum samples obtained from n=587 participants in Replication Cohort II.

| **Inflammatory marker** | **Overall (n=1,170)** | | **Trimester 1* (n=421)** | | **Trimester 2* (n=347)** | | **Trimester 3* (n=402)** | | **p-value**** |
| --- | --- | --- | --- | --- | --- | --- | --- | --- | --- |
|  | **Median (IQR)** | **Range (min-max)** | **Median (IQR)** | **Range (min-max)** | **Median (IQR)** | **Range (min-max)** | **Median (IQR)** | **Range (min-max)** |  |
| IL-1β (pg/ml) | 2.14 (1.33) | 0.26-82.96 | 2.18 (1.48) | 0.26-82.96 | 2.14 (1.35) | 0.43-9.39 | 2.12 (1.26) | 0.48-48.96 | 0.180 |
| IL-6 (pg/ml) | 1.79 (1.74) | 0.03-109.28 | 1.94 (1.97) | 0.10-109.28 | 1.76 (1.60) | 0.03-86.92 | 1.75 (1.69) | 0.15-72.76 | 0.150 |
| IL-17A (pg/ml) | 16.55 (10.88) | 1.48-160.56 | 17.3 (11.14) | 2.24-96.58 | 16.05 (10.35) | 1.48-84.8- | 16.09 (9.97) | 2.41-160.56 | 0.088 |
| HS-CRP (mg/L) | 4.78 (4.04) | 0.23-55.33 | 3.27 (4.1) | 0.24-44.48 | 3.70 (3.92) | 0.30-43.89 | 3.22 (3.96) | 0.23-55.33 | 0.426 |

**Timepoint according to study design: Timepoint 1 = median 12 weeks gestation. Timepoint 2 = median 20 weeks gestation. Timepoint 3 = median 28 weeks gestation.

**One-way ANOVA of normalized cytokines and HS-CRP between trimesters.

**Supplementary Table 3: Proportion of variance in inflammatory markers explained by prominent drivers.** Variance partitioning analysis showed that individual was the main driver of inflammatory markers in the Discovery Cohort, Replication Cohort I and Replication Cohort II. Only drivers explaining more than 1% are depicted in the table.

|  | **Model Without CRP PGS** | | | **Model With CRP PGS** | | | |
| --- | --- | --- | --- | --- | --- | --- | --- |
|  | **Individual (%)** | **BMI**  **(%)** | **Residuals (%)** | **Individual (%)** | **BMI**  **(%)** | **PGS all (%)** | **Residuals (%)** |
| *Discovery Cohort* | | | |  |  |  |  |
| HS-CRP | 55.6 | 9.6 | 32.1 | 41.1 | 9.4 | 14.1 | 32.7 |
| IL-1β | 82.2 | 0.2 | 16.3 | 82.8 | 0.3 | 0.0 | 14.4 |
| IL-6 | 77.0 | 0.1 | 21.9 | 78.4 | 0.1 | 0.0 | 20.4 |
| IL-17A | 86.2 | 0.5 | 11.7 | 85.2 | 0.5 | 0.0 | 12.0 |
| IL-23 | 92.8 | 0.1 | 6.6 | 92.9 | 0.0 | 0.0 | 6.6 |
| IFN-γ | 86.8 | 0.4 | 10.9 | 86.5 | 0.3 | 0.0 | 11.1 |
| Maternal Cytokine Index | 87.4 | 0.2 | 10.5 | 86.6 | 0.2 | 0.0 | 10.4 |
| *Replication Cohort I* | | | |  |  |  |  |
| HS-CRP | 33.4 | 14.9 | 50.5 |  |  |  |  |
| IL-1β | 26.3 | 0.0 | 72.9 |  |  |  |  |
| IL-6 | 71.5 | 0.0 | 28.0 |  |  |  |  |
| IL-17A | 68.2 | 0.6 | 29.6 |  |  |  |  |
| *Replication Cohort II* | | | |  |  |  |  |
| HS-CRP | 58.1 | 16.1 | 25.1 |  |  |  |  |
| IL-1β | 80.9 | 0.1 | 18.7 |  |  |  |  |
| IL-6 | 74.7 | 0.1 | 24.5 |  |  |  |  |
| IL-17A | 84.2 | 0.5 | 14.7 |  |  |  |  |

**Supplementary Table 4: Mixed effects model to investigate the association between selected predictors and HS-CRP in Replication Cohort I and II.**

|  | HS-CRP  Replication Cohort I (n=3,319) | | | | HS-CRP  Replication Cohort II (n=1,170) | | | |
| --- | --- | --- | --- | --- | --- | --- | --- | --- |
|  | $\beta$ | 95% CI | *P*-value | Corrected P-value* | $\beta$ | 95% CI | *P*-value | Corrected P-value* |
| Maternal age | 0.00 | -0.01; 0.01 | 0.484 | 0.610 | 0.00 | -0.03; 0.02 | 0.763 | 0.763 |
| National Background** | -0.14 | -0.24; -0.05 | **0.002** | **0.005** | 0.15 | -0.16; 0.46 | 0.347 | 0.623 |
| Pre-pregnancy BMI | 0.07 | 0.06; 0.07 | **<0.001** | **0.004** | 0.14 | 0.11; 0.16 | **<0.001** | **0.009** |
| Unemployment | NA | NA | NA |  | 0.09 | -0.37; 0.55 | 0.698 | 0.763 |
| Multiparity | 0.08 | 0.00; 0.17 | 0.059 | 0.094 | 0.16 | -0.04; 0.36 | 0.126 | 0.378 |
| Maternal tobacco use  During pregnancy | NA | NA | NA | NA | 0.29 | -0.48; 1.06 | 0.463 | 0.695 |
| Maternal alcohol use  During pregnancy | NA | NA | NA | NA | -0.18 | -1.21; 0.85 | 0.737 | 0.763 |
| Female fetus pregnancy | -0.02 | -0.10; 0.06 | 0.610 | 0.610 | -0.12 | -0.31; 0.08 | 0.233 | 0.524 |
| Twin pregnancy | 0.08 | -0.22; 0.38 | 0.600 | 0.610 | NA | NA | NA | NA |
| Private insurance | -0.10 | -0.21; 0.00 | 0.051 | 0.094 | NA | NA | NA | NA |
| Gestational age of the sample | 0.01 | 0.00; 0.01 | **0.001** | **0.004** | 0.01 | 0.00; 0.02 | **0.019** | 0.086 |
| Marginal R^2^ / Conditional R^2^ | 0.178/0.506 | | | | 0.187/0.745 | | | |

*p-value after Benjamini-Hochberg correction.

** National Background indicates White ethnicity for Replication Cohort I, and Dutch national background for Replication Cohort II.

N/A indicates that data was specific to the Discovery Cohort and was not measured in the replication cohorts.

**Supplementary Table 5: Characteristics of inflammatory markers in the pre-pregnancy cohort (Generation R *Next* cohort).** Inflammatory marker levels overall, as well as per timepoint in maternal serum samples obtained from n=1,270 participants.

| **Inflammatory marker** | **Overall**  **(n=1,779)** | | **Timepoint 1***  **(n=676)** | | **Timepoint 2***  **(n=1,103)** | | **p-value**** |
| --- | --- | --- | --- | --- | --- | --- | --- |
|  | **Median (IQR)** | **Range (min-max)** | **Median (IQR)** | **Range (min-max)** | **Median (IQR)** | **Range (min-max)** |  |
| IL-1β (pg/ml) | 4.82 (3.35) | 0.72-174.18 | 4.82 (3.18) | 0.77-174.18 | 4.84 (3.44) | 0.72-159.13 | 0.474 |
| IL-6 (pg/ml) | 1.92 (1.93) | 0.02-74.17 | 2.00 (2.17) | 0.02-48.32 | 1.89 (1.79) | 0.04-74.17 | 0.248 |
| IL-17A (pg/ml) | 59.47 (35.62) | 8.44-1,447.29 | 60.65 (35.45) | 11.66-1447.29 | 58.52 (35.94) | 8.44-707.36 | 0.840 |
| IL-23 (pg/ml) | 1,209.5 (882.02) | 128.7-41,122.54 | 1,199.3 (926.21) | 166.50-30,690.10 | 1230.00 (858.53) | 128.70-41,122.50 | 0.323 |
| IFN-*y* (pg/ml) | 18.89 (11.74) | 1.39-365.81 | 19.13 (11.15) | 1.39-365.81 | 18.72 (11.96) | 1.39-103.50 | 0.130 |
| HS-CRP (mg/L) | 1.40 (2.90) | 0.60-74.00 | 0.90 (1.70) | 0.60-36.00 | 11.80 (3.30) | 0.60-74.00 | **0.008** |

*Timepoint according to study design: Timepoint 1 = preconception, Timepoint 2= first trimester, median 8.4 weeks gestation.

**Paired samples t-test of normalized cytokines and HS-CRP preconception and in the first trimester.

**Supplementary Table 6: Mixed effects models to investigate the association between selected predictors and HS-CRP and the maternal cytokine index in a sensitivity analysis excluding samples with potential HAAA interference (n=163 samples)**

|  | HS-CRP (n=9,889) | | | | Maternal cytokine index^+^ (n=9,994) | | | |
| --- | --- | --- | --- | --- | --- | --- | --- | --- |
|  | $\beta$ | 95% CI | *P*-value | Corrected P-value* | $\beta$ | 95% CI | *P*-value | Corrected P-value* |
| Maternal age | 0.00 | 0.00 – 0.01 | 0.336 | 0.448 | -0.01 | -0.02 – 0.00 | **<0.001** | **0.004** |
| Dutch national background | 0.00 | -0.06 – 0.06 | 0.959 | 0.959 | 0.08 | 0.03 – 0.13 | **0.003** | **0.008** |
| Pre-pregnancy BMI | 0.09 | 0.08 – 0.09 | **<0.001** | **0.003** | -0.01 | -0.02 – 0.00 | **0.001** | **0.004** |
| CRP PGS (all ancestry) | 0.45 | 0.42 – 0.48 | **<0.001** | **0.003** | -0.01 | -0.03 – 0.02 | 0.578 | 0.771 |
| PGS (European ancestry)*** | 0.47 | 0.44 – 0.50 | **<0.001** | **0.003** | 0.00 | -0.03 – 0.03 | 0.978 | 0.997 |
| High income | -0.12 | -0.18 – -0.05 | **<0.001** | **0.003** | 0.04 | -0.02 – 0.09 | 0.217 | 0.362 |
| Multiparity | 0.23 | 0.18 – 0.29 | **<0.001** | **0.003** | -0.03 | -0.08 – 0.02 | 0.261 | 0.402 |
| Maternal Psychopathology | 0.01 | -0.07 – 0.08 | 0.851 | 0.896 | 0.01 | -0.05 – 0.08 | 0.656 | 0.772 |
| Maternal tobacco use**  Pre-pregnancy | -0.02 | -0.12 – 0.07 | 0.620 | 0.729 | -0.13 | -0.22 – -0.05 | **0.002** | **0.007** |
| Maternal tobacco use **  During pregnancy | 0.05 | -0.02 – 0.12 | 0.195 | 0.300 | -0.18 | -0.24 – -0.11 | **<0.001** | **0.004** |
| Maternal alcohol use**  Pre-pregnancy | -0.07 | -0.16 – 0.01 | 0.092 | 0.153 | 0.02 | -0.06 – 0.10 | 0.617 | 0.771 |
| Maternal alcohol use**  Occasionally during pregnancy | -0.11 | -0.17 – -0.04 | **0.001** | **0.003** | 0.05 | 0.00 – 0.11 | 0.067 | 0.134 |
| Maternal alcohol use**  Frequently during pregnancy | -0.22 | -0.33 – -0.11 | **<0.001** | **0.003** | 0.02 | -0.08 – 0.12 | 0.702 | 0.780 |
| Substance use during pregnancy | -0.13 | -0.24 – -0.02 | **0.019** | **0.035** | 0.04 | -0.05 – 0.14 | 0.384 | 0.549 |
| Female fetus pregnancy | -0.02 | -0.07 – 0.03 | 0.424 | 0.530 | 0.00 | -0.05 – 0.05 | 0.997 | 0.997 |
| Twin pregnancy | -0.05 | -0.31 – 0.22 | 0.721 | 0.801 | 0.19 | -0.04 – 0.43 | 0.111 | 0.202 |
| Season | 0.03 | 0.01 – 0.04 | **0.008** | **0.018** | -0.02 | -0.03 – -0.01 | **<0.001** | **0.004** |
| Infection score*** | 0.04 | 0.01 – 0.06 | **0.001** | **0.003** | 0.02 | 0.01 – 0.03 | **0.003** | **0.008** |
| Gestational age of the sample | -0.01 | -0.01 – 0.00 | **0.010** | **0.020** | -0.01 | -0.02 – -0.01 | **<0.001** | **0.004** |
| Cytokine batch | 0.00 | 0.00 – 0.00 | 0.267 | 0.381 | 0.00 | 0.00 – 0.00 | **0.018** | **0.040** |
| Marginal R^2^ / Conditional R^2^ | 0.252/0.672 | | | | 0.018/0.886 | | | |

+ Maternal cytokine index was scaled

*p-value after Benjamini-Hochberg correction.

** Reference group: Never smoked during pregnancy/Never drank during pregnancy

***Interpreted in separate models.

**Supplementary Table 7: Mixed effects models to investigate the association between selected predictors and HS-CRP and the maternal cytokine index in a sensitivity analysis excluding samples of participants with auto-immune diseases (n=855 samples).**

|  | HS-CRP (n=9,208) | | | | Maternal cytokine index^+^ (n=9,302) | | | |
| --- | --- | --- | --- | --- | --- | --- | --- | --- |
|  | $\beta$ | 95% CI | *P*-value | Corrected P-value* | $\beta$ | 95% CI | *P*-value | Corrected P-value* |
| Maternal age | 0.00 | 0.00 – 0.01 | 0.182 | 0.274 | -0.01 | -0.02 – -0.01 | **<0.001** | **0.007** |
| Dutch national background | 0.00 | -0.06 – 0.06 | 0.991 | 0.991 | 0.09 | 0.03 – 0.14 | **0.004** | **0.016** |
| Pre-pregnancy BMI | 0.09 | 0.08 – 0.09 | **<0.001** | **0.003** | -0.01 | -0.01 – 0.00 | **0.011** | **0.028** |
| PGS (all ancestry) | 0.46 | 0.43 – 0.48 | **<0.001** | **0.003** | 0.00 | -0.03 – 0.02 | 0.901 | 0.901 |
| PGS (European ancestry)*** | 0.48 | 0.45 – 0.51 | **<0.001** | **0.003** | 0.01 | -0.02 – 0.04 | 0.600 | 0.800 |
| High income | -0.12 | -0.19 – -0.05 | **0.001** | **0.003** | 0.03 | -0.03 – 0.10 | 0.314 | 0.523 |
| Multiparity | 0.23 | 0.18 – 0.29 | **<0.001** | **0.003** | -0.02 | -0.07 – 0.04 | 0.519 | 0.741 |
| Maternal Psychopathology | 0.01 | -0.07 – 0.09 | 0.850 | 0.895 | 0.01 | -0.07 – 0.08 | 0.848 | 0.893 |
| Maternal tobacco use**  Pre-pregnancy | -0.02 | -0.12 – 0.08 | 0.739 | 0.821 | -0.13 | -0.23 – -0.04 | **0.005** | **0.017** |
| Maternal tobacco use **  During pregnancy | 0.05 | -0.03 – 0.12 | 0.221 | 0.295 | -0.18 | -0.26 – -0.11 | **<0.001** | **0.007** |
| Maternal alcohol use**  Pre-pregnancy | -0.06 | -0.15 – 0.03 | 0.162 | 0.270 | 0.03 | -0.05 – 0.12 | 0.452 | 0.695 |
| Maternal alcohol use**  Occasionally during pregnancy | -0.11 | -0.18 – -0.04 | **0.002** | **0.006** | 0.06 | 0.00 – 0.13 | 0.051 | 0.093 |
| Maternal alcohol use**  Frequently during pregnancy | -0.23 | -0.34 – -0.11 | **<0.001** | **0.003** | 0.02 | -0.08 – 0.13 | 0.643 | 0.804 |
| Substance use during pregnancy | -0.12 | -0.23 – 0.00 | **0.043** | 0.078 | 0.01 | -0.09 – 0.12 | 0.796 | 0.893 |
| Female fetus pregnancy | -0.03 | -0.08 – 0.03 | 0.303 | 0.356 | -0.01 | -0.06 – 0.05 | 0.836 | 0.893 |
| Twin pregnancy | -0.15 | -0.43 – 0.12 | 0.267 | 0.334 | 0.26 | 0.01 – 0.51 | **0.0**4**5** | 0.090 |
| Season | 0.03 | 0.01 – 0.04 | **0.009** | **0.020** | -0.02 | -0.03 – -0.01 | **0.002** | **0.010** |
| Infection score*** | 0.04 | 0.01 – 0.06 | **0.003** | **0.008** | 0.02 | 0.01 – 0.03 | **0.007** | **0.020** |
| Gestational age of the sample | -0.01 | -0.01 – 0.00 | **0.028** | 0.056 | -0.01 | -0.02 – -0.01 | **<0.001** | **0.007** |
| Cytokine batch | 0.00 | 0.00 – 0.00 | 0.192 | 0.274 | 0.00 | 0.00 – 0.00 | **0.039** | 0.087 |
| Marginal R^2^ / Conditional R^2^ | 0.256/0.669 | | | | 0.016/0.890 | | | |

+ Maternal cytokine index was scaled

*p-value after Benjamini-Hochberg correction.

** Reference group: Never smoked during pregnancy/Never drank during pregnancy

***Interpreted in separate models.

**Supplementary Table 8: Mixed effects models to investigate the association between selected predictors and HS-CRP and the maternal cytokine index in a sensitivity analysis excluding outlier samples (excluding n=174 samples).**

|  | HS-CRP (n=9,874) | | | | Maternal cytokine index^+^ (n=9,874) | | | |
| --- | --- | --- | --- | --- | --- | --- | --- | --- |
|  | $\beta$ | 95% CI | *P*-value | Corrected P-value* | $\beta$ | 95% CI | *P*-value | Corrected P-value* |
| Maternal age | 0.00 | 0.00 – 0.01 | 0.381 | 0.508 | -0.01 | -0.02 – -0.01 | **<0.001** | **0.003** |
| Dutch national background | -0.01 | -0.07 – 0.05 | 0.795 | 0.837 | 0.07 | 0.02 – 0.12 | **0.012** | **0.030** |
| Pre-pregnancy BMI | 0.09 | 0.08 – 0.09 | **<0.001** | **0.003** | -0.01 | -0.02 – 0.00 | **0.001** | **0.003** |
| PGS (all ancestry) | 0.44 | 0.42 – 0.47 | **<0.001** | **0.003** | 0.00 | -0.03 – 0.02 | 0.811 | 0.901 |
| PGS (European ancestry)*** | 0.46 | 0.43 – 0.49 | **<0.001** | **0.003** | 0.00 | -0.03 – 0.03 | 0.965 | 0.965 |
| High income | -0.10 | -0.17 – -0.04 | **0.002** | **0.004** | 0.03 | -0.03 – 0.09 | 0.272 | 0.453 |
| Multiparity | 0.22 | 0.17 – 0.28 | **<0.001** | **0.003** | -0.02 | -0.07 – 0.03 | 0.391 | 0.569 |
| Maternal Psychopathology | 0.02 | -0.05 – 0.09 | 0.582 | 0.667 | -0.01 | -0.08 – 0.05 | 0.711 | 0.873 |
| Maternal tobacco use**  Pre-pregnancy | 0.01 | -0.09 – 0.10 | 0.906 | 0.906 | -0.14 | -0.23 – -0.06 | **0.001** | **0.003** |
| Maternal tobacco use **  During pregnancy | 0.06 | -0.01 – 0.13 | 0.115 | 0.177 | -0.17 | -0.24 – -0.11 | **<0.001** | **0.003** |
| Maternal alcohol use**  Pre-pregnancy | -0.07 | -0.16 – 0.01 | 0.081 | 0.135 | 0.01 | -0.06 – 0.09 | 0.742 | 0.873 |
| Maternal alcohol use**  Occasionally during pregnancy | -0.11 | -0.17 – -0.04 | **0.001** | **0.003** | 0.07 | 0.01 – 0.13 | **0.022** | **0.049** |
| Maternal alcohol use**  Frequently during pregnancy | -0.21 | -0.32 – -0.11 | **<0.001** | **0.003** | 0.03 | -0.07 – 0.12 | 0.606 | 0.808 |
| Substance use during pregnancy | -0.14 | -0.25 – -0.04 | **0.007** | **0.014** | 0.04 | -0.05 – 0.14 | 0.398 | 0.569 |
| Female fetus pregnancy | -0.01 | -0.07 – 0.04 | 0.588 | 0.667 | 0.00 | -0.04 – 0.05 | 0.880 | 0.926 |
| Twin pregnancy | -0.07 | -0.32 – 0.18 | 0.600 | 0.667 | 0.24 | 0.01 – 0.47 | **0.044** | 0.088 |
| Season | 0.02 | 0.00 – 0.03 | 0.067 | 0.122 | -0.02 | -0.03 – -0.01 | **<0.001** | **0.003** |
| Infection score*** | 0.04 | 0.02 – 0.06 | **<0.001** | **0.003** | 0.01 | 0.00 – 0.02 | 0.146 | 0.265 |
| Gestational age of the sample | -0.01 | -0.01 – 0.00 | **0.001** | **0.003** | -0.01 | -0.02 – -0.01 | **<0.001** | **0.003** |
| Cytokine batch | 0.00 | 0.00 – 0.00 | 0.209 | 0.299 | 0.00 | 0.00 – 0.00 | **0.011** | **0.030** |
| Marginal R^2^ / Conditional R^2^ | 0.259/0.689 | | | | 0.018/0.899 | | | |

+ Maternal cytokine index was scaled

*p-value after Benjamini-Hochberg correction.

** Reference group: Never smoked during pregnancy/Never drank during pregnancy

***Interpreted in separate models.

**Supplementary Table 9: Mixed effects models to investigate the association between selected predictors and HS-CRP and the maternal cytokine index in a subgroup of participants with a high absolute HS-CRP delta (5^th^ and 95^th^ quantile of the HS-CRP delta between timepoints) (n=854 samples)**

|  | HS-CRP (n=854) | | | | Maternal cytokine index^+^ (n=854) | | | |
| --- | --- | --- | --- | --- | --- | --- | --- | --- |
|  | $\beta$ | 95% CI | *P*-value | Corrected P-value* | $\beta$ | 95% CI | *P*-value | Corrected P-value* |
| Maternal age | 0.01 | -0.02 – 0.04 | 0.622 | 0.822 | -0.02 | -0.04 – 0.00 | 0.107 | 0.668 |
| Dutch national background | 0.04 | -0.26 – 0.35 | 0.779 | 0.822 | 0.12 | -0.10 – 0.34 | 0.290 | 0.778 |
| Pre-pregnancy BMI | 0.05 | 0.01 – 0.09 | **0.010** | 0.060 | -0.02 | -0.04 – 0.01 | 0.258 | 0.778 |
| CRP PGS (all ancestry) | 0.46 | 0.33 – 0.58 | **<0.001** | **0.010** | -0.04 | -0.13 – 0.05 | 0.425 | 0.869 |
| CRP PGS (European ancestry)*** | 0.50 | 0.35 – 0.64 | **<0.001** | **0.010** | -0.03 | -0.13 – 0.08 | 0.599 | 0.929 |
| High income | -0.33 | -0.65 – -0.01 | **0.040** | 0.133 | 0.02 | -0.21 – 0.25 | 0.844 | 0.929 |
| Multiparity | 0.16 | -0.11 – 0.43 | 0.242 | 0.605 | 0.14 | -0.06 – 0.33 | 0.167 | 0.668 |
| Maternal Psychopathology | 0.08 | -0.29 – 0.46 | 0.663 | 0.822 | 0.01 | -0.26 – 0.29 | 0.929 | 0.929 |
| Maternal tobacco use**  Pre-pregnancy | 0.22 | -0.29 – 0.73 | 0.405 | 0.816 | -0.04 | -0.41 – 0.33 | 0.828 | 0.929 |
| Maternal tobacco use **  During pregnancy | 0.02 | -0.38 – 0.42 | 0.929 | 0.929 | -0.22 | -0.51 – 0.07 | 0.134 | 0.668 |
| Maternal alcohol use**  Pre-pregnancy | -0.11 | -0.54 – 0.31 | 0.605 | 0.822 | -0.05 | -0.36 – 0.25 | 0.732 | 0.929 |
| Maternal alcohol use**  Occasionally during pregnancy | -0.11 | -0.41 – 0.20 | 0.500 | 0.822 | -0.06 | -0.28 – 0.16 | 0.605 | 0.929 |
| Maternal alcohol use**  Frequently during pregnancy | -0.08 | -0.58 – 0.42 | 0.749 | 0.822 | -0.02 | -0.39 – 0.35 | 0.904 | 0.929 |
| Substance use during pregnancy | -0.17 | -0.78 – 0.44 | 0.580 | 0.822 | 0.08 | -0.36 – 0.52 | 0.729 | 0.929 |
| Female fetus pregnancy | -0.09 | -0.35 – 0.16 | 0.473 | 0.822 | -0.07 | -0.25 – 0.12 | 0.478 | 0.869 |
| Twin pregnancy | -0.19 | -1.52 – 1.14 | 0.781 | 0.822 | 0.78 | -0.20 – 1.57 | 0.120 | 0.668 |
| Season | 0.10 | -0.01 – 0.20 | 0.072 | 0.206 | 0.02 | -0.02 – 0.05 | 0.311 | 0.778 |
| Infection score*** | 0.16 | 0.03 – 0.28 | **0.013** | 0.060 | 0.01 | -0.04 – 0.05 | 0.802 | 0.929 |
| Gestational age of the sample | 0.04 | 0.01 – 0.08 | **0.015** | 0.060 | -0.01 | -0.02 – 0.00 | **0.007** | 0.140 |
| Cytokine batch | -0.01 | -0.02 – 0.01 | 0.408 | 0.816 | 0.00 | 0.00 – 0.01 | 0.442 | 0.869 |
| Marginal R^2^ / Conditional R^2^ | 0.088/0.673 | | | | 0.024/0.880 | | | |

+ Maternal cytokine index was scaled

*p-value after Benjamini-Hochberg correction.

** Reference group: Never smoked during pregnancy/Never drank during pregnancy

***Interpreted in separate models.

**Supplementary Table 10: Immunoassay performance (n=13,744)**. Detection of each analyte in the immunoassay was >99.5%. Less than 0.1% of the samples interfered with the analyte’s beads resulting in low bead count and these were excluded. Samples that fell outside the range of the standard curve (out of range) were substituted with the lowest value for each analyte. After exclusion and substitution, n=13,467 samples were included in the current study.

| **Analyte** | **Detection** | **Low bead count, n (%)** | **N OOR, n (%)** | **Substituted value for OOR samples (pg/mL)** |
| --- | --- | --- | --- | --- |
| IL-1β | 99.9% | 3, <0.1% | 9, <0.1% | 0.28 |
| IL-6 | 99.6% | 3, <0.1% | 57, 0.4% | 0.01 |
| IL-17A | 99.9% | 2, <0.1% | 5, <0.1% | 1.05 |
| IL-23 | 99.9% | 3, <0.1% | 7, <0.1% | 63.48 |
| IFN-γ | 99.9% | 4, <0.1% | 9, <0.1% | 0.9 |

Legend: OOR = out of range.

**Supplementary Table 11: Characteristics of inflammatory markers prior to and after exclusion of samples with potential human anti-animal antibodies (HAAA) interference in the Discovery Cohort.** Samples with potential HAAA interference were excluded in sensitivity analyses. Median inflammatory marker levels were significantly higher when potential HAAA samples were included.

| **Inflammatory marker** | **Including potential HAAA samples (n=13,467)** | | **Excluding potential HAAA samples (n=13,255)** | | ***p*-value** |
| --- | --- | --- | --- | --- | --- |
|  | **Median (IQR)** | **Range (min/max)** | **Median (IQR)** | **Range (min/max)** |  |
| IL-1β (pg/ml) | 3.95 (2.0) | 0.12 – 5195.84 | 3.92(1.89) | 0.12 – 5,195.84 | **<0.01** |
| IL-6 (pg/ml) | 1.61 (1.57) | 0.00 – 507.61 | 1.6 (1.46) | 0.01 – 168.64 | **<0.01** |
| IL-17A (pg/ml) | 23.91 (16.46) | 0.50 – 2493.46 | 23.62 (15.57) | 0.5 – 433.41 | **<0.01** |
| IL-23 (pg/ml) | 1102.50 (1019.25) | 29.22 – 143087.31 | 1,088.31 (972.6) | 29.22 – 119,986.29 | **<0.01** |
| IFN-γ (pg/ml) | 14.75 (10.71) | 0.71 – 999.42 | 14.53 (10.31) | 0.71 –315.28 | **<0.01** |
|  | **Including potential HAAA samples (n=13,316)*** | | **Excluding potential HAAA samples (n=13,110)*** | |  |
| HS-CRP (mg/L) | 4.3 (5.0) | 0.2 – 343.0 | 4.3 (5.1) | 0.2 – 343.0 | 0.838 |

*HS-CRP was measured in 13,316 samples (5,924 early and 7,395 mid pregnancy samples).

**Supplementary Table 12: Validation of the Polygenic Score (PGS) of CRP.** The PGS of CRP was validated by assessing the change in R-squared upon adding it to the null model of maternal serum HS-CRP (log2 normalized). The significant increase in R-squared validates the PGS.

|  | R^2^ |
| --- | --- |
| Null Model^*^ | 0.02324 |
| Null Model^*^ + PGS | 0.16404 |
| *R^2^ Change* | *0.14080* |

^*^ Null Model includes PC 1 to PC 10 as predictors and HS-CRP (log2 normalized) as outcome.
